# *Orientia tsutsugamushi:* analysis of the mobilome of a highly fragmented and repetitive genome reveals ongoing lateral gene transfer in an obligate intracellular bacterium

**DOI:** 10.1101/2023.05.11.540415

**Authors:** Suparat Giengkam, Chitrasak Kullapanich, Jantana Wongsantichon, Haley E. Adcox, Joseph J. Gillespie, Jeanne Salje

**Affiliations:** Mahidol-Oxford Tropical Medicine Research Unit, Faculty of Tropical Medicine, Mahidol University, Bangkok, Thailand; Department of Microbiology and Immunology, Virginia Commonwealth University Medical Center, School of Medicine, Richmond, Virginia, USA; Department of Microbiology and Immunology, School of Medicine, University of Maryland Baltimore, MD 21201; Department of Pathology, Department of Biochemistry, Cambridge Institute for Medical Research, University of Cambridge, UK; Public Health Research Institute, Rutgers the State University of New Jersey, Newark, USA

**Keywords:** *Orientia tsutsugamushi*, Rickettsiales, obligate intracellular bacteria, intracellular pathogens, mobile genetic elements, comparative genomics, bacteriophage, integrative and conjugative elements, lateral gene transfer.

## Abstract

The rickettsial human pathogen *Orientia tsutsugamushi* (Ot) is an obligate intracellular Gram-negative bacterium with one of the most highly fragmented and repetitive genomes of any organism. Around 50% of its ∼2.3 Mb genome is comprised of repetitive DNA that is derived from the highly proliferated Rickettsiales amplified genetic element (RAGE). RAGE is an integrative and conjugative element (ICE) that is present in a single Ot genome in up to 92 copies, most of which are partially or heavily degraded. In this report, we analysed RAGEs in eight fully sequenced Ot genomes and manually curated and reannotated all RAGE-associated genes, including those encoding DNA mobilisation proteins, P-type (*vir*) and F-type (*tra)* type IV secretion system (T4SS) components, Ankyrin repeat- and tetratricopeptide repeat-containing effectors, and other piggybacking cargo. Originally, the heavily degraded Ot RAGEs led to speculation that they are remnants of historical ICEs that are no longer active. Our analysis, however, identified two Ot genomes harbouring one or more intact RAGEs with complete F-T4SS genes essential for mediating ICE DNA transfer. As similar ICEs have been identified in unrelated rickettsial species, we assert that RAGEs play an ongoing role in lateral gene transfer within the Rickettsiales. Remarkably, we also identified in several Ot genomes remnants of prophages with no similarity to other rickettsial prophages. Together these findings indicate that, despite their obligate intracellular lifestyle and host range restricted to mites, rodents and humans, Ot genomes are highly dynamic and shaped through ongoing invasions by mobile genetic elements and viruses.

## Importance

Obligate intracellular bacteria, or those only capable of growth inside other living cells, have limited opportunities for horizontal gene transfer with other microbes due to their isolated replicative niche. The human pathogen *Orientia tsutsugamushi* (Ot), an obligate intracellular bacterium causing scrub typhus, encodes an unusually high copy number of a ∼40 gene mobile genetic element that typically facilitates genetic transfer across microbes. This proliferated element is heavily degraded in Ot and previously assumed to be inactive. Here, we conducted detailed analysis of this element in eight Ot strains and discovered two strains with at least one intact copy. This implies that the element is still capable of moving across Ot populations and suggests that the genome of this bacterium may be even more dynamic than previously appreciated. Our work raises questions about intracellular microbial evolution and sounds an alarm for gene-based efforts focused on diagnosing and combatting scrub typhus.

## Introduction

***Orientia tsutsugamushi*** (**Ot**) is an obligate intracellular Gram-negative bacterium that is a symbiont of trombiculid mites and causes the vector-borne human disease scrub typhus. Ot is a member of the alphaproteobacterial order Rickettsiales, which contains three well-studied families: Anaplasmataceae, Rickettsiaceae and Midichloriaceae^1,2^, as well as four lesser-known families that have recently been described (Deianiraeaceae, Mitibacteraceae, Gamibacteraceae, and Athabascaceae)^3–5^. As a lineage within Rickettsiaceae, genus *Orientia* also includes “*Candidatus* Orientia chiloensis”, which has recently been identified as an endemic species in Chile^6^, and Candidatus *O. chuto*, which was isolated from a patient in Dubai^7^. There is extensive strain diversity within the Ot species, which can be found in rodents, mites and human patients across Southeast Asia. While strain diversity corresponds to differences in virulence in patients and in animal infection models, the molecular basis of these differences in virulence are not well understood. Ot strains are often classified according to serotype groupings, which are organised based on the human serological response to the highly antigenic surface protein TSA56. Major serotype groups are named after type strains and include Karp, Kato, Gilliam, Japanese-Gilliam, TA763, Saitama, Kuroki, Kawasaki and Shimokoshi.

At around 2-2.5 Mb, the genome of Ot is almost double the size of most Rickettsiales genomes and is one of the most fragmented and repetitive bacterial genomes reported to date^8, 9^. With almost 50% of the genome comprised of repetitive DNA sequences, many experimental approaches are challenging: i.e. primer design, gene and genome sequencing, gene prediction and annotation, and comparative genomics. Complete genome sequences of two strains, Boryong and Ikeda, were published in 2008 using short read sequencing and bacterial artificial chromosome cloning^8, 9^. An additional six strains (Karp, Kato, Gilliam, TA686, UT76, UT176) were fully sequenced in 2018 using long read PacBio technology^10^. A comparison of these eight genomes enabled the identification of 657 core genes and an open pangenome that is heavily characterized by gene duplication and pseudogenisation rather than the import of novel genes^10^.

The Ot genome is dominated by an **integrative and conjugative element** (**ICE**), called **Rickettsiales amplified genetic element** (**RAGE**)^8, 9, 11^, that has proliferated rampantly throughout the genome and is present in over 70 copies. RAGEs encode numerous repeated and pseudogenized genes, as well as single copy cargo genes that appear to be important for bacterial growth and pathogenesis. Whilst the high number of RAGE copies in the Ot genome remains unmatched, similar RAGEs have been described in several other *Rickettsia* species: *Rickettsia bellii* (single intact copy^12^), *Rickettsia buchneri* (7 complete or near-complete genomic copies, and two plasmid encoded copies)^11^, *Rickettsia massiliensis* (single intact copy^13^), *R*. *parkeri* str. Atlantic Rainforest (single intact copy^14^), *R. felis* str. LSU-Lb (one plasmid encoded copy^15^) and *R. peacockii* (one partially degraded copy)^16^. In Ot, the ICE has not been controlled by the bacterial host and the RAGE has replicated to high levels in the genome^9, 10, 17^. The reasons for the differential fate of the RAGEs in Rickettsiaceae are unknown, but the small effective population size as well as the presence of population bottlenecks in obligate intracellular bacteria likely explain why it has had the ability to proliferate without strong negative selection in at least some rickettsial species.

Here, we present a thorough Ot phylogenomics analysis and re-annotation of the RAGEs and their cargo genes in eight strains: Gilliam, Boryong, UT76, UT176, Karp, Kato, Ikeda and TA686. We delineate the start and stop sites of all the intact and degraded RAGEs, allowing us to identify **inter-RAGE** (**IR**) regions with conserved clusters of genes. We further describe complete RAGEs in two Ot genomes (Kato and Gilliam). Finally, we annotate and classify the genes associated with DNA mobilisation and divergent **type IV secretion systems** (**T4SSs**), as well as the numerous multicopy cargo genes within the RAGEs, including those encoding **Ankyrin-repeat containing proteins** (**Anks**) and **tetratricopeptide repeat containing proteins** (**TPRs**), which are putative secreted effectors. This detailed disentangling of the superfluous RAGE-dominated mobilome from the core and accessory Ot genome is expected to enlighten research on Ot biology and overall genome evolution in obligate intracellular bacteria.

## Results and discussion

### RAGE and IR regions

The RAGE is an ICE that is present in Ot and certain other Rickettsiales genomes. The degree of amplification and degradation of RAGE in the Ot genome is so extensive - making up around 50% of the Ot genome in 71-93 distinct genomic regions – that the beginning and end sites of RAGEs cannot be easily identified by visual inspection. Here, we established objective criteria for the classification of genes into Ot RAGEs, and manually delineated each RAGE in the eight Ot strains in our study.

First, one or more copies of each of the following mobilisation genes must be present: integrase (*int)*, transposase (*tnp)*, F-type T4SS genes (*tra/trb)*, and relaxosome (*tra*). Second, one or more previously defined cargo or regulation genes^9^ must be present: membrane proteins (reclassified here as Ot_RAGE_membrane protein, see below), DNA adenine methyltransferase (*dam*), DNA helicase, ATP-binding proteins (*mrp*), histidine kinases, SpoT-related proteins (synthetase and/or hydrolase domains), HNH endonuclease, peroxiredoxin, Anks, and TPRs. Third, the RAGE region begins with the first RAGE-associated gene and continues until a previously defined core gene^10^ is reached. A run of core genes is classified as an IR element. Fourth, given the abundance of genes encoding **hypothetical proteins** (**HPs**) within RAGEs, those located between mobilisation/cargo RAGE genes are classified as being part of that RAGE. However, HP-encoding genes located between RAGE mobilisation/cargo genes and Ot core genes that cannot be resolved as being within RAGE or IR regions are classified as isolated HP-encoding genes. Fifth and final, single or multiple mobilisation or cargo RAGE genes are classified as isolated mobile genes or cargo genes, respectively. Some new cargo genes were identified by virtue of residing within RAGE regions in most or all genomes and these are discussed below.

In this way the entire genome of each Ot strain was classified into the following regions: RAGE region, IR region, isolated HP-encoding gene, isolated mobilisation gene, and isolated cargo gene (**Supplementary Dataset 1**). Using these criteria, we identified 71-93 RAGEs in the eight analysed genomes (**Fig. 1A, B**). The patterns of RAGE fragmentation and pseudogenization varied extensively between strains and it was not possible to map RAGEs between strains (**Fig. 1A**). This implies that RAGEs entered Ot strains one or more times as intact elements and subsequently underwent replication, pseudogenisation and recombination in independent trajectories.

**Figure 1.**
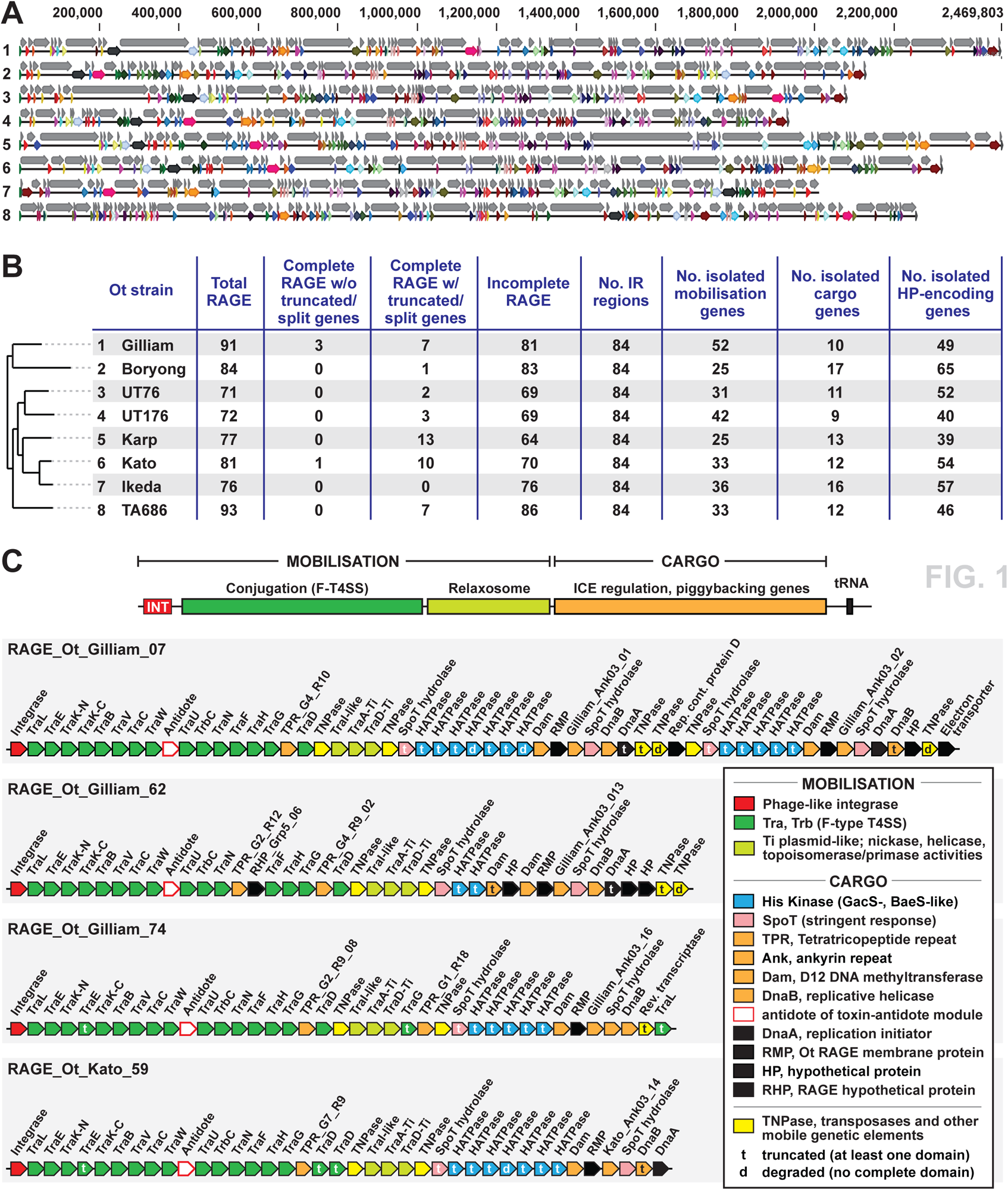
Ot RAGE and IR elements. **A.** An overview of the genomes of eight Ot strains with genes classified into RAGE and IR regions. Numbers at left refer to Ot strains listed in panel B. Grey arrows = RAGE regions; colored arrows = IR regions. The colors correspond to conserved IR regions between strains and demonstrate the lack of synteny between Ot genomes. **B.** Table summarizing RAGEs, IR regions and isolated mobile genes, cargo genes and hypothetical genes that could not be classified into RAGE or IR elements. Ot strains are listed accordingly to a previously estimated phylogeny^10^, with numbers corresponding to full genome maps in panel A. **C.** Organization of genes in the four complete RAGEs found in our analysis. t = truncated (at least one identifiable domain present); d = degraded (no identifiable domains present). Detailed analysis of the reannotation and classification of all genes in the eight genomes are given in **Supplementary Dataset 1**.

Most RAGEs in Ot are degraded, both in terms of either completely lacking RAGE-associated genes or retaining genes that have been truncated or fragmented into predicted pseudogenes. A previous study found that the Ot strain Ikeda genome lacked any complete RAGES^9^. We assessed whether any of the strains in our analysis encoded complete RAGEs, defined as containing a full set of mobilisation genes and additional cargo genes as outlined in previous analyses^9^. All the strains in our analysis, with the notable single exception of Ikeda, encoded one or more complete set of RAGE genes (**Fig. 1B**). However, most of those RAGEs contained one or more mobilisation genes that were truncated. Accordingly, we carried out sequence alignments to define each RAGE gene as being full-length, truncated (containing one or more identifiable domains) or degraded (containing no identifiable complete domains). We then assessed whether any strains contained complete RAGEs with intact, full-length genes (**Fig. 1B**). We found that two strains, Gilliam and Kato, encoded complete RAGEs with full length mobilisation genes (**Fig. 1C**). This suggests that these strains may have obtained these elements recently and that they may be capable of mobilisation. ICEs normally have a preferred integration site, often within tRNA genes^11^, as observed for *Rickettsia* species^18^, Despite our discovery of complete RAGEs in these Ot genomes, no identifiable integration sites could be determined.

We previously used RNA sequencing analysis and comparative genomics to show that, despite the lack of synteny between Ot strains driven by the prolific RAGEs, small groups of proximate genes were transcribed at similar levels and maintained synteny across strains^19^. This demonstrates selection for gene order at the local level despite it being absent at a global level across Ot genomes. To further identify gene groups evolving under strong selective constraints relative to superfluous RAGEs, we analysed the IR regions, which harbour the majority of core Ot genes (**Fig. 1A**). Remarkably, this revealed 84 IR regions, ranging in length from 2 to 27 genes, most of which were conserved across all strains (**Fig. 1B**). Identification of these conserved IR gene groups illuminates highly conserved microsynteny that may encompass functionally linked genes sharing expression and/or regulatory programs.

### Single copy cargo genes

The delineation of Ot genomes into RAGE and IR regions enabled us to better characterize RAGE cargo genes (**Fig. 2A, B**). In addition to the group of highly replicated multicopy cargo genes already described as RAGE components (discussed below), we identified numerous single copy genes previously overlooked for their occurrence within RAGEs (**Fig. 2A**). These include genes involved in fundamental processes of bacterial physiology and metabolism, e.g., tyrosine tRNA ligase (*tyrS),* RNA polymerase subunit omega (*rpoZ),* and the ClpP protease (*clpP)*, as well as genes encoding predicted secretory effectors likely involved in interactions with host cells, including phospholipase D (*pld)* and autotransporter proteins ScaA and ScaC (*scaA, scaC*) in all genomes, and ScaB (*scaB,* Boryong), ScaF (*scaF,* TA686) and ScaG (*scaG,* TA686) in individual strains. As many of these single copy genes have orthologs in other bacterial species that lack the RAGEs, it is likely that they were not introduced by mobile genetic elements. Rather, their current presence within RAGEs indicates they were probably incorporated into RAGE via recombination. However, a case-by-case basis may reveal certain conserved genes shuttling between Ot genomes via RAGE mobilisation. For instance, despite their conservation in all *Rickettsia* genomes, genes encoding secreted effectors and metabolite transporters were previously found piggybacking on RAGEs in the *R*. *buchneri* genome, illustrating the ability for RAGE to shuttle rickettsial genes important for the obligate intracellular lifestyle^11^.

**Figure 2.**
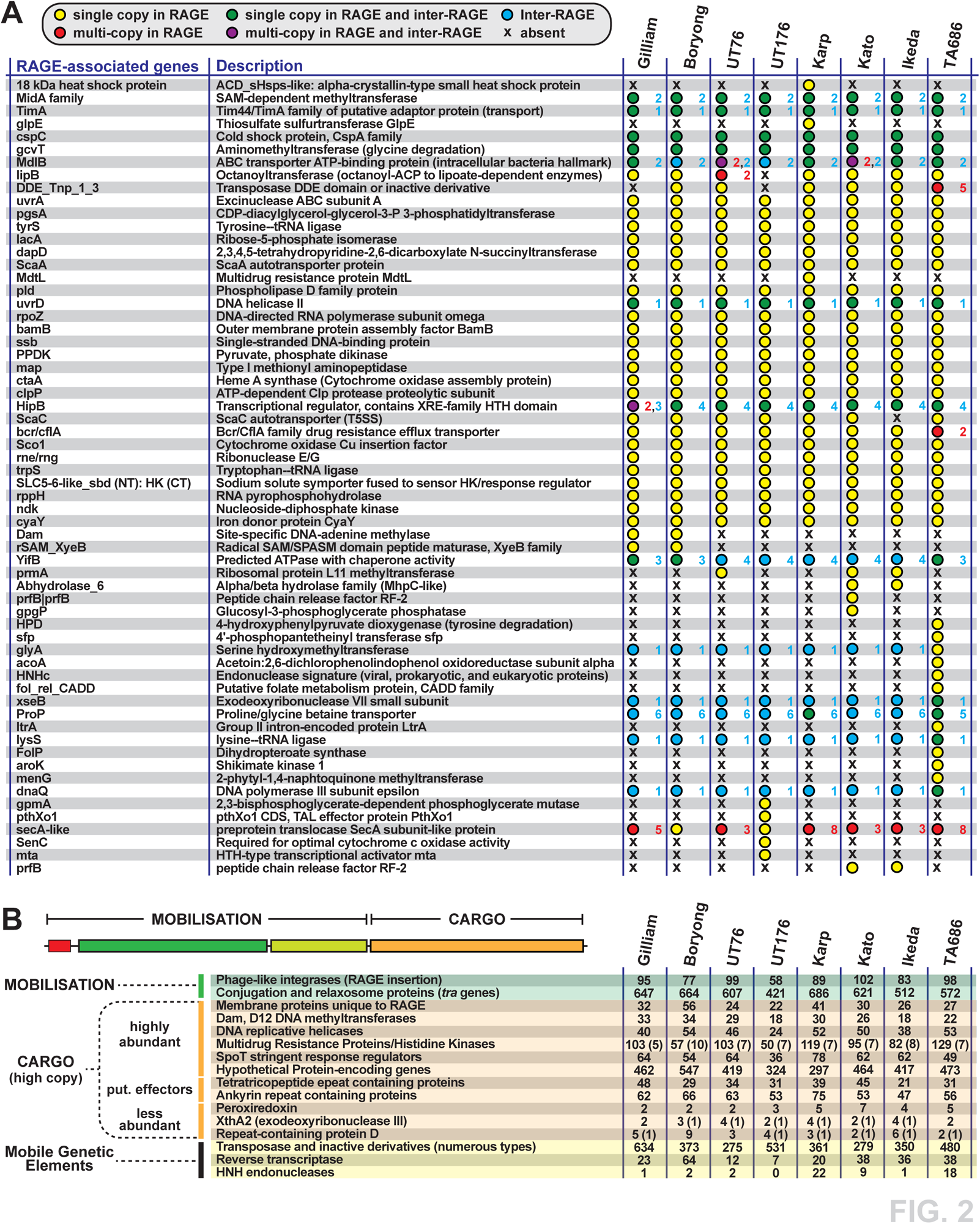
Single and multi-copy cargo genes encoded on Ot RAGEs. **A.** Single or low-copy cargo genes encoded on Ot RAGE. Summary statistics show whether genes are present in single or multiple copies on RAGEs in different strains, and also in single or multiple copies in IRs. The exact number of copies is given for each gene. Blue text = number of copies in IR; red text = number of copies in RAGE. **B.** Frequency and distribution of high copy cargo genes (both full length and truncated/degraded) within RAGEs in eight strains of Ot. Numbers in brackets denote additional copies in IRs.

### Highly abundant multi-copy cargo genes

Analysis of RAGE-associated cargo genes revealed 16 genes or (gene groups) present in numerous copies in all eight Ot genomes (**Fig. 2B**). Gene groups included membrane proteins, Dam DNA methyltransferases, DNA helicases, **multidrug resistance proteins** (**MRP**) and histidine kinases, SpoT hydrolase and synthetases, hypothetical/uncharacterized genes, mobile genetic elements (i.e., insertion sequences, transposases, integrases, and reverse transcriptases), Anks, TPRs, and *vir*- and *tra-*type T4SS genes. All of these, except *vir-*type T4SS genes, have been identified as RAGE cargo genes in previous studies^8, 9, 20, 21^. For each gene group we assessed (i) whether all the genes annotated as belonging to this category were paralogs of the same gene or whether multiple distinct genes were present within one group, and (ii) whether some or all genes within a group were truncated and not able to form a full-length protein and, where a functional domain was known, whether this domain was present or not. Genes involved in DNA mobilisation, effector proteins, and T4SS genes are discussed in dedicated sections below, whilst other multi-copy cargo genes are discussed here.

#### Membrane proteins

The eight Ot genomes encode 21-41 RAGE associated genes annotated as membrane proteins (**Fig. 3A, Supplementary Dataset 2**). Analyses revealed that each Ot strain encodes exactly one copy of three genes encoding proteins with analogy to characterized membrane proteins: the YccA modulator of protease FtsH, vitamin transporter Vut1 and a gene similar to the rhamnose transporter RhaT. The remaining 18-38 genes encode paralogs of a gene we call Ot_RAGE_membrane protein, ranging in length from 90 to 663 bp. This protein lacks homology to any non-Ot genes and no known domain could be identified. Thus, the function of this gene in Ot is unknown.

**Fig. 3.**
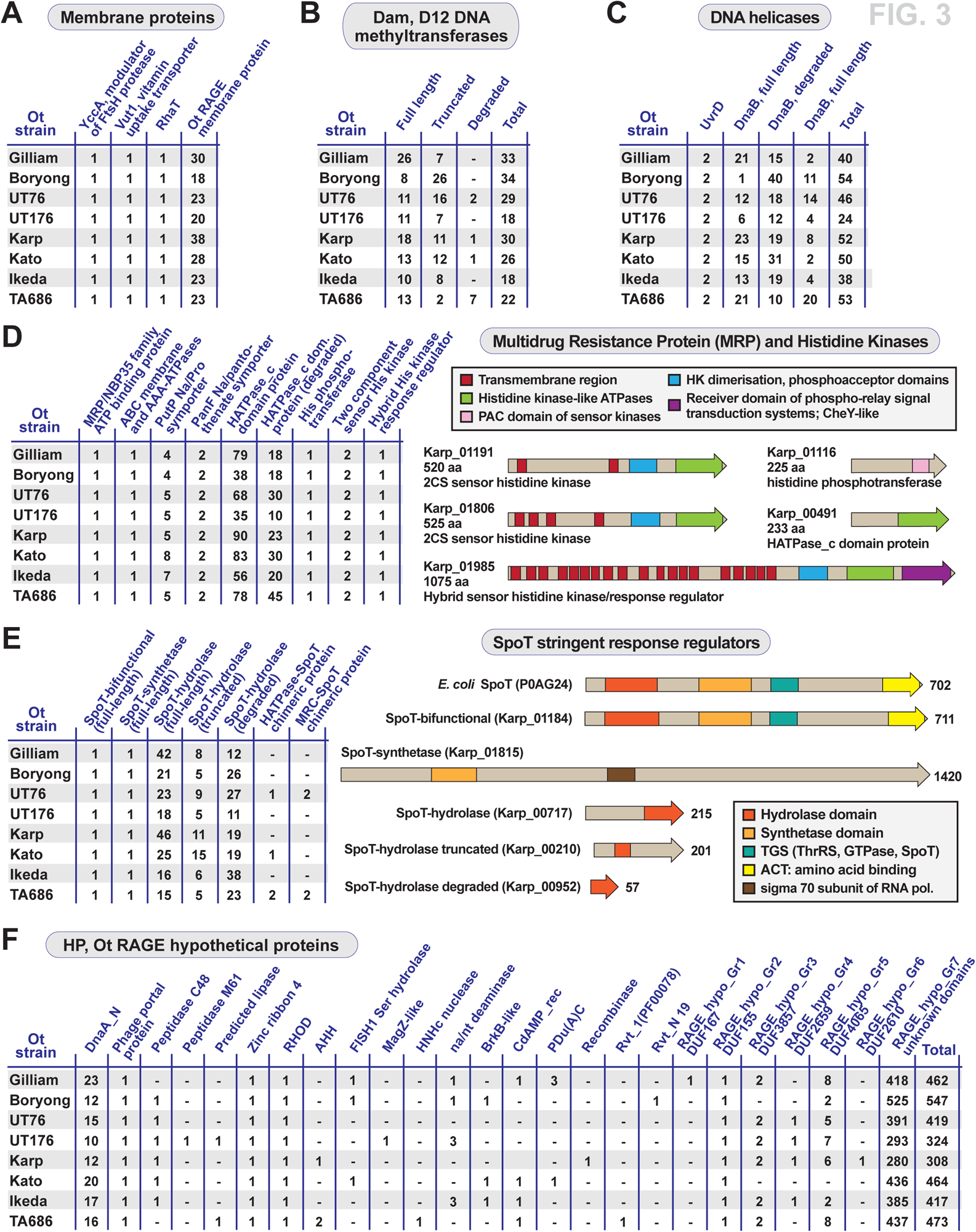
Analysis of high copy cargo genes on RAGE elements in Ot. **A-C.** Frequency and distribution of RAGE cargo genes annotated as (**A**) membrane proteins, (**B**) Dam DNA methyltransferases, and (**C**) DNA helicases. DnaB is a replicative DNA helicase and UvrB is a repair DNA helicase. **D**. Frequency and distribution of RAGE cargo genes encoding MRP/histidine kinases, with examples of His kinase divergent architectures. **E**. Frequency and distribution of RAGE cargo genes encoding SpoT stringent response regulators, with examples of divergent architectures. The bifunctional SpoT protein is compared to the canonical SpoT protein of *E*. *coli*. **E**. Frequency and distribution of RAGE cargo genes encoding HPs. DnaA_N, N-terminal domain of DnaA; RHOD, rhodanese homology domain; AHH, adenosyl homocysteine hydrolase; MagZ, nucleoside triphosphate pyrophospho-hydrolase; na/nt, nucleic acid/nucleotide deaminase; BrkB-like, YihY/virulence factor BrkB family protein; PDu(A)C, copper chaperone; CdAMP_rec, cyclic diAMP receptor proteins; Rvt_1 (PF00078), reverse transcriptase Pfam PF00078; Rvt_N 19, domain of reverse transcriptase Rvt_N; DUF, domain of unknown function.

#### DNA methyltransferases

Ot genomes encode 18 to 34 genes with similarity to DNA adenine methyltransferase (*dam*), all of which are located within RAGES (**Fig. 3B**). Sequence alignments demonstrate that 8-26 are full length proteins, defined as being equal in length to *E. coli dam* and encoding all seven known domains. The Ot genomes encode an additional 2-26 truncated *dam* genes where some domains are preserved, and fewer degraded copies with no identifiable domains. It is not known if these genes are functional, although previous studies^19, 22^ showed that they were not detected by proteomics analysis. They may have a specific role in methylation of RAGE during mobilisation and/or integration to protect from deleterious effects of single-stranded DNAse activity.

#### DNA helicases

DNA helicases unwind double stranded DNA and function in DNA and RNA metabolism, with general roles in DNA replication, repair and recombination. We found that all strains of Ot encode exactly two full length copies of the DNA helicase UvrD, which is involved in DNA repair (**Fig. 3C**). One copy is located within a RAGE whilst the second is located within IR82. The DnaB family of helicases, by contrast, is more numerous and degraded in Ot genomes (**Fig. 3C**). This helicase is involved in DNA replication and is present in 36-52 copies, with 1-23 being full length and mostly located within RAGEs. Each genome encodes one full length copy located at the interface of IR46 and IR47, which is likely the ancestral non-RAGE paralog involved in genome replication. Other copies are undoubtedly associated with RAGE mobilisation. Our previous proteomics analysis^19, 22^ detected expression of a DnaB gene, but due to sequence similarities between numerous paralogs, it was not possible to determine which specific gene(s) was expressed. Given its role in DNA replication, however, it is expected that at least one paralog would be expressed and functional.

#### MRPs and histidine kinases

We identified around 100 genes in the Ot genomes that were annotated as MRPs or histidine kinases (**Fig. 2B**, **3D**, **Supp. Fig. 1, Supplementary Dataset 3)**, or were found to contain histidine kinase domains (e.g. the sodium/proline symporter PutP). MRPs are members of the **ATP-binding cassette** (**ABC**) transporter protein family and were annotated as MRPs based on the presence of a **histidine kinase ATPase domain** (**HATPase**). Our analysis determined that two “MRP” proteins were distinct from all the other HATPase domain containing proteins in Ot: an MRP/NBP35 family ATP-binding protein and an ABC-membrane and AAA ATPase protein, both single copy in each genome. The former was located within an IR region (IR9) and the latter within RAGEs. Analysis of the remaining genes annotated as MRP/histidine kinases led us to identify several full-length orthologs of **two-component system** (**2CS**) histidine kinase genes. These include one histidine phosphotransferase gene (which does not contain an HATPase domain), one large hybrid sensor histidine kinase/response regulator gene and two 2CS histidine kinase genes (**Fig. 3D**). All were present in the same copy number and locations in each genome. Histidine phosphotransferase and the two 2CS sensor histidine kinase genes were consistently found in IR regions IR43, IR47, and IR63, respectively, whilst the hybrid sensor histidine kinase/response regulator gene was located within a RAGE. In our UT76 proteomics dataset^22^, both the histidine phosphotransferase and the hybrid sensor histidine kinase/response regulator genes were detected, whilst the two sensor histidine kinase proteins were not (**Supplementary Dataset 1**). We also identified two copies of a histidine kinase domain containing sodium/pantothenate symporter, PanF, present in IR11 and IR55 regions in each genome, as well as a sodium/proline symporter, PutP, present in 4-8 copies and distributed into both RAGEs and IRs. Analysis of our previous proteomics dataset showed that PanF was expressed in strain UT76 as was one copy of PutP located in IR49^22^ (**Supplementary Dataset 1**). The remaining MRP/histidine kinase genes were paralogs of one another, containing an HATPase domain and being present in 45-113 copies per genome with various degrees of truncation (**Supp. Fig. 1**). We classified these as degraded HATPase domain containing proteins when an intact HATPase domain could no longer be detected due to the short length of the gene (**Fig. 3D, Supp. Fig. 1, Supplementary Dataset 1, 3**).

#### SpoT stringent response regulators

SpoT is a bifunctional synthetase/hydrolase that is essential for inducing and regulating the stringent response in *E. coli* and other bacteria through mediating intercellular levels of alarmone, or (p)ppGpp^23, 24^. Ot genomes encode 36-78 genes with homology to SpoT. We identified exactly one full length SpoT gene, present in an IR region (IR46), encoding both synthetase and hydrolase domains in all Ot genomes (**Fig. 3E**). This gene was the only SpoT homolog found to be expressed in our previous proteomics analysis and was shown to be upregulated in extracellular Ot, consistent with a role in transitioning between different bacterial states^22^. We also identified exactly one gene in each genome that encodes only the SpoT synthetase domain in addition to a long C-terminal domain of unknown function. We then identified 15-46 SpoT genes that lacked the synthetase domain yet encoded the intact hydrolase domain as well as a further 16-44 SpoT genes that were truncated or degraded such that a functional hydrolase domain was no longer present.. Finally, we identified 1-4 genes in some genomes in which hydrolase domains are fused to other genes. Most rickettsial genomes harbour 6-12 SpoT genes, with some of the abovementioned architectures present (data not shown). Curiously, bifunctional (complete hydrolase and synthetase domains) genes are typical of most other Rickettsiales species, though not common in *Rickettsia* species and absent from notable human pathogens (e.g., *R*. *prowazekii*, *R*. *typhi*, *R*. *rickettsii*, and *R*. *conorii*)^11^. Still, the tendency for all Rickettsiales genomes to retain numerous single domain SpoT genes, even when RAGEs are absent, implies their function in some aspect of the stringent response. The presence of such drastic numbers and diverse architectures of SpoT genes in Ot genomes relative to other rickettsial species is intriguing and deserving of future investigation.

#### HPs

The Ot strains harbour 308 to 547 genes per genome that are annotated as hypothetical or uncharacterized, of which about half are located within RAGEs (**Fig. 2B** and **3F and Supplementary Dataset 4**). We determined whether all the RAGE hypothetical/uncharacterized genes were paralogs of a single RAGE gene or if they encoded multiple different genes. Sequence alignments for all the genes annotated as hypothetical or uncharacterized in the Karp genome were performed, which revealed that the genes clustered into 24 groups (**Fig. 3F**) with 18 of these encoding genes carrying known protein domains. Those without known domains were named RAGE_hypo_Gr1-7, with groups 1-6 encoding a domain of unknown function, and group 7 combining all remaining HP genes with no identifiable known domains. Three of these genes with known domains were present in exactly one copy in all genomes and encode: a phage portal protein, a zinc ribbon domain, and a rhodanese homology domain. Another single-copy gene found in all eight Ot genomes carries a domain of unknown function (DUF155). Furthermore, hypothetical genes containing a DnaA N-terminal domain were identified in 10-23 copies in all Ot genomes. In our prior study, one full length paralog, located in IR1 was expressed in UT76, whilst others were not detected^22^ (**Supplementary Dataset 1**).

Other hypothetical/uncharacterized genes were distributed sporadically amongst the genomes. In order to get a sense of the distribution of the remaining RAGE-associated hypothetical genes that were not clustered into conserved groups, we analysed the remaining hypothetical genes in the RAGEs of the Karp genome only. There were no identifiable domains in any of these and the diversity was such that it was not possible to bin them into homologous groups. We annotated them all as belonging to a large and divergent 25^th^ group (RAGE_hypo_Gr7). While little can be inferred about the function of these hundreds of genes, it is likely that at least some of these play important roles in the biology of Ot.

### Putative effectors piggybacking on Ot RAGE

#### Anks

The ankyrin repeat is one of the most common protein folds in nature, being widespread in eukaryotes and pervasive in many viruses and host-associated bacteria^25, 26^. Ankyrin repeats are used to mediate a myriad of protein-protein interactions, and host-associated prokaryotes and viruses frequently express Anks to hijack or subvert host cell pathways that would be detrimental or beneficial to their survival^27^. Previously, several Ot Anks were shown to be secreted via the rickettsial **type I secretion system** (**T1SS**)^28^. Certain Ank effectors have been functionally characterized in strain Ikeda and shown to play important roles in host cell interactions^28–32^. However, a major challenge in comparing the host-pathogen cell biology of different Ot strains has been the difficulty assessing which Anks are most similar to those in other strains. This is important for determining the significance of Anks as species-versus strain-specific effectors underlying pathogenesis.

We defined a set of criteria for clustering Anks, with their subsequent characterisation within each Ot genome following the well described Ank repertoire of strain Ikeda ^9^. We identified several new Ank groups in Ikeda, although some of these lack complete Ank repeats and are likely non-functional **Supplementary Dataset 5**). Our comparative analysis indicates Ot strains encode 47-66 Anks, with variability (67-94%) in the number of common vs. strain-specific proteins per genome (**Fig. 4A**). Ot Anks often harbour a single F-box domain, which are prominently known components of SCF (Skp1, Cullin1, F-box) ubiquitin ligase complexes but recent studies have described their participation in non-SCF protein-protein interactions involved in diverse eukaryotic functions?pathways?^33^. F-box-resembling PRANC (pox protein repeats of ankyrin-C-terminal) domains and coiled-coils were less frequently predicted.

**Figure 4.**
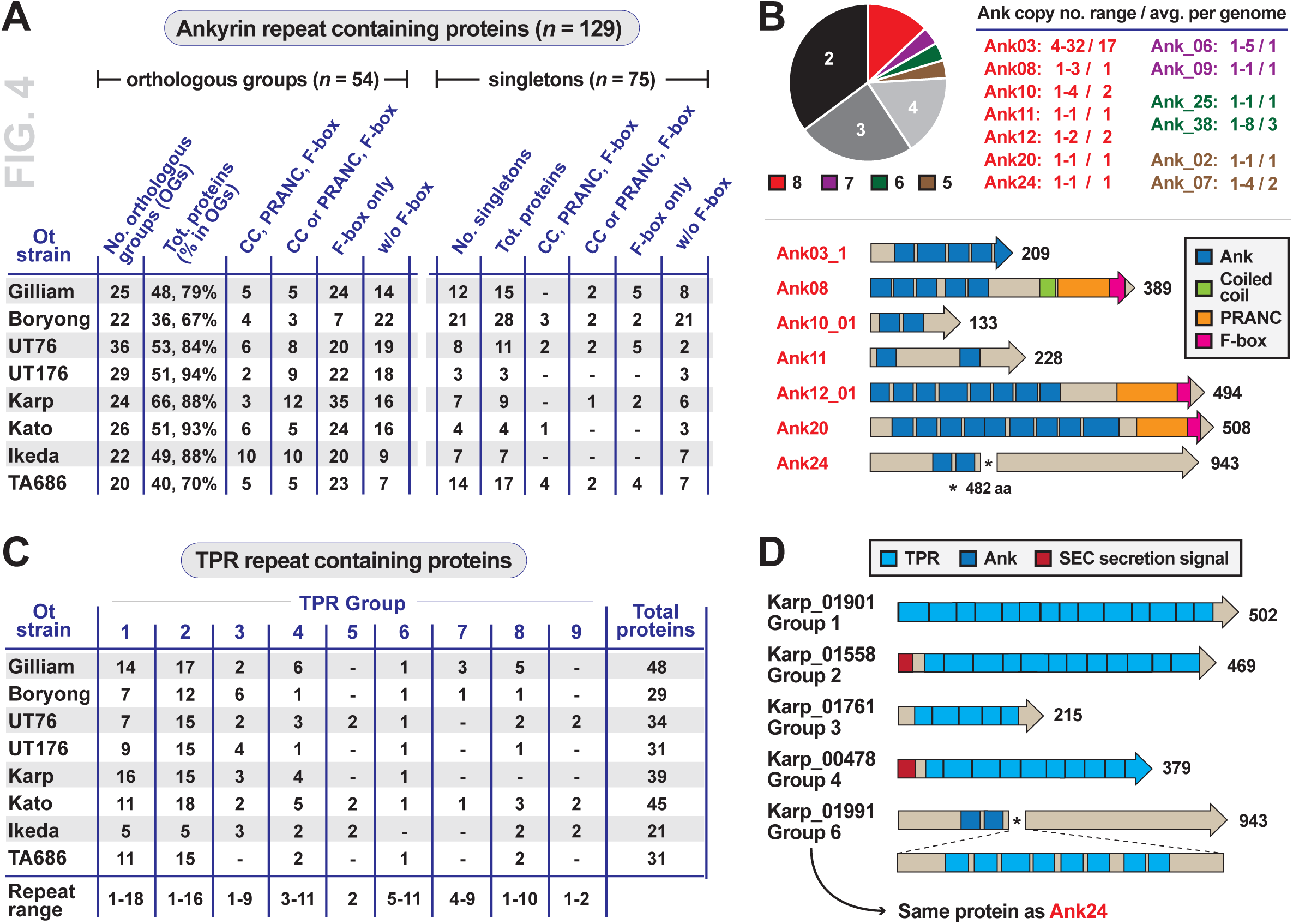
Putative effectors piggybacking on Ot RAGE. **A**. Frequency and distribution of Anks in Ot genomes. Anks are broken down into orthologous groups (OGs, present in two or more genomes) or singletons (unique to a genome). CC, coiled coil; PRANC (Pox proteins Repeats of ANkyrin, C-terminal), domain found at the C terminus of certain Pox virus proteins; F-box, motif of approximately 50 amino acids that functions in protein-protein interactions. **B**. (*top*) graphical view of Ank OG strain representation (2-8 genomes). Roughly 25% of Ank OGs are found in five or more strains, with variable levels of conservation in copy number per genome. (*bottom*) Architectures for Anks present in all Ot genomes, with proteins from Ot strain Karp. **C.** Frequency and distribution of TPRs in Ot genomes. Nine ortholog groups contain all the TPR s across eight Ot genomes. **D.** Examples of diverse TPR architectures for six proteins from Ot strain Karp.

Of the 54 orthologous groups of Ot Anks, all genomes were found to encode at least one copy of seven groups: Ank03, Ank08, Ank10, Ank11, Ank12, Ank20 and Ank24 (**Fig. 4B**). Ank03 is by far the most prominent Ank, being present in 4 to 32 copies in Ot genomes, with the other six families present in 1 to 4 copies (**Fig. 4B**). Curiously, while most Ot Anks are predominantly found within RAGEs, Ank20 is encoded in an IR region (IR84) in all analysed Ot genomes. Collectively, these seven Anks likely carry out essential functions in Ot biology. However, each strain likely utilizes unique Ank arsenals throughout its lifecycle given that some of the less conserved Anks have characterized functional roles; e.g, Ank01 and Ank06 of Ot str. Ikeda modulate NFkB transport to the nucleus^31^. As such, it is likely that there is functional redundancy between the Ank groups, with some of the 100 other Ank groups not found in Ikeda functioning similarly as Ank01 and Ank06 in genomes lacking these genes.

#### TPRs

The tetratricopeptide repeat is another protein motif that is commonly used in mediating inter-protein interactions, typically found in subunits of multi-protein complexes^34^. TPRs are widespread in Ot proteins (**Fig. 4C**), albeit with a lower number of copies per genome than Anks. Ot TPRs have been less characterised than the Ot Anks, with only one report demonstrating a role in inhibition of eukaryotic translation in Ot strain Boryong^35^. We compiled 21-48 TPRs per Ot genome and classified them into nine groups primarily based on within-protein location of tetratricopeptide repeats (**Fig. 4C, Supp. Fig. 2, Supplementary Dataset 5**). Whilst the positions were conserved within groups, the number of repeats was variable and indicated expansion and contraction of repeats, as well as processive gene degradation within each group (**Fig. 4D**). The prediction of SEC signal peptides in certain TPRs indicates at least some of these putative effectors may be secreted to the periplasm with possible translocation across the outer membrane, possibly via TolC as proposed for the RARP-1 effector of *R*. *typhi*^36^. Still, the lack of N-terminal secretion signals in most TPRs indicates other possible routes for TPR secretion that await characterisation.

### Mobile genetic elements associated with RAGE

#### Integrases

ICEs, such as Ot RAGE, encode integrase genes to catalyse genomic integration, and conjugative genes (discussed in the next section) to catalyse horizontal gene transfer^37^. The Ot genomes analysed in this study encode 58-102 integrase genes (**Fig. 5A**), of which only 3-13 per genome remain full length, consistent with progressive degradation of the Ot RAGE. Where present, the integrases are located at the start position of a RAGE (**Fig. 1C**). However, several integrase genes were located as isolated genes outside RAGE regions, reflecting the high mobility of these genes and the overall high recombination rates in Ot genomes.

**Figure 5.**
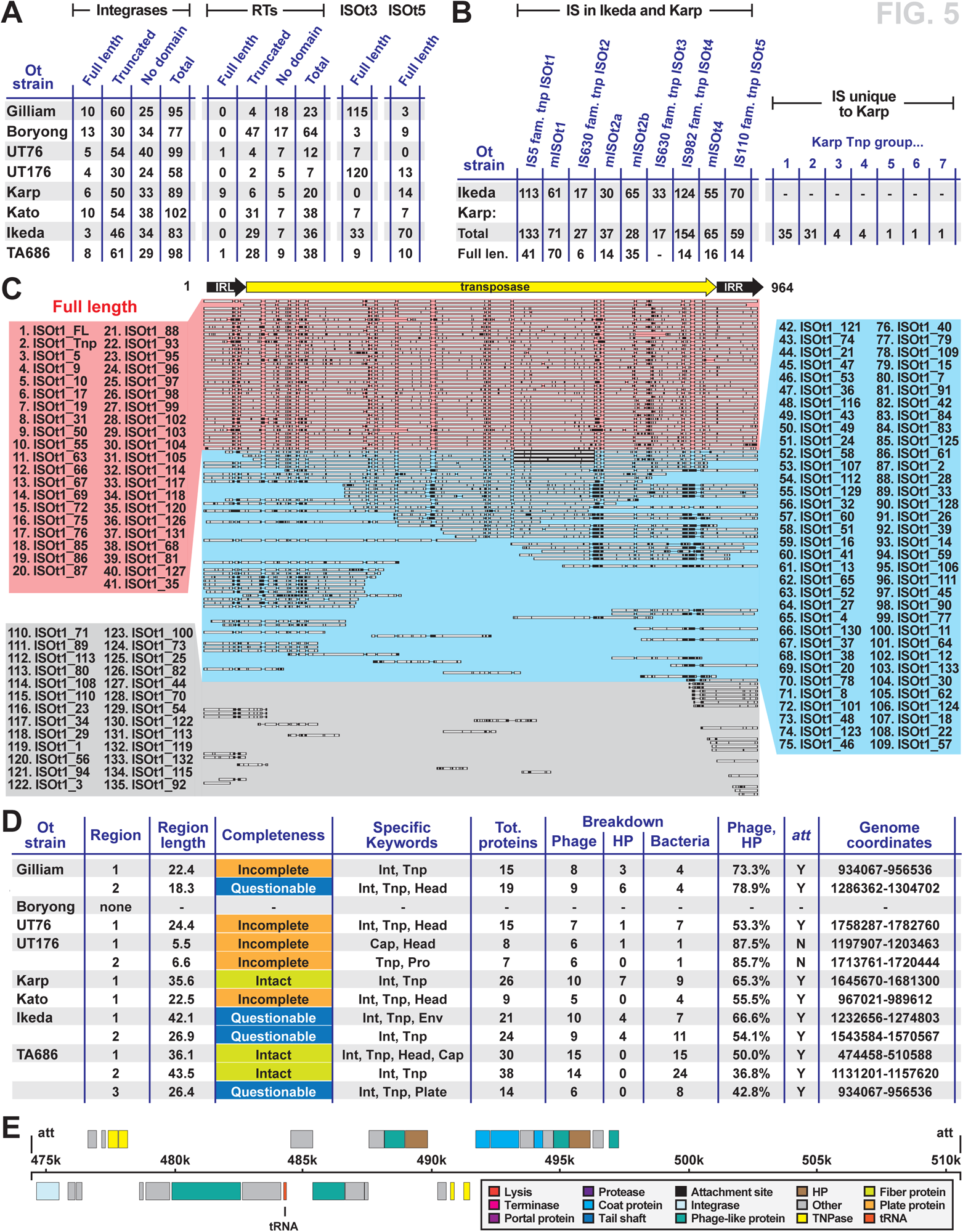
A diversity of mobile genetic elements associates with Ot RAGEs. **A**. Frequency and distribution of Ot RAGE-associated genes encoding integrases, Group II intron-associated reverse transcriptases, and IS elements ISOt3 and ISOt5. **B**. Frequency and distribution of IS elements in Ikeda and Karp strains. **C**. Alignment showing classification of ISOt1 elements as full length or degraded. Full length copies of ISOt1 in Karp are shown by red dotted box. **D**. Overview of prophage elements in Ot genomes as identified by PHASTER search tool. Int = integrase, Tnp = transposase, Env = envelope, Cap = capsid, Pro = protease. **E**. Overview of predicted phage region in TA686.

#### Transposable elements

In addition to the OtRAGE, the Ot genome encodes two other types of transposable elements^9^: retrotransposons (group II introns) and DNA transposons. Whilst these are independent mobile genetic elements, they have been incorporated into the Ot RAGE regions, and it is likely that the different mobilizable elements impact each other’s activity. Group II introns are self-splicing retrotransposons that catalyse their integration into genomes via an RNA intermediate, using an intron-encoded reverse transcriptase protein^38^. The Ot genomes encode 7-64 group II intron reverse transcriptase genes, although only UT76, Karp and TA686 encode full length genes, with the others being heavily degraded (**Fig. 5A**). All full-length reverse transcriptase genes were immediately followed by an HNH endonuclease gene likely required for catalysis.

Ot encodes several families of DNA transposons which have been previously classified in strain Ikeda^9^. This class of mobile elements is comprised of a transposase gene flanked by inverted repeat regions on either side, which together make up an **insertion sequence** (**IS**)^39^. Numerous families of IS have been identified in other bacteria. Given the large number of IS elements in each Ot genome and their highly degrative tendency, we selected two of the many frequently occurring IS genes, ISOt3 and ISOt5, to characterise in detail across all eight genomes (**Fig. 5A**). These were present in 0-120 (ISOt3) and 0-70 (ISOt5) full-length copies across the genomes. We also analysed the complete set of IS elements in one strain, Karp, and compared these with those previously predicted for str. Ikeda (**Fig. 5B**). An example of the analysis of one IS element in Karp, ISOt1, shows the distribution of full length and degraded copies typical of IS families in all Ot genomes (**Fig. 5C**). We identified the same set of IS elements that had previously been described in Ikeda^9^ (**Fig. 5B, C**). We followed the classification and nomenclature established in Nakayaka et al^9^, in which mISOt1, mISOt2 and mISOt4 denotes “miniature” versions of elements containing the same terminal inverted repeat sequences as ISOt1, ISOt2 and ISOt4 respectively. Within the nine IS classes found in Karp, most were heavily degraded with some, such as IS630 family transposase ISOt3, having no remaining full-length elements. We identified an additional seven groups of transposase genes in Karp that were not part of IS elements (**Fig. 5B**).

Bacteriophages are another source of horizontally transferred genetic material. These are thought to be rare in obligate intracellular bacteria due to the isolated lifestyle, although there are exceptions such as the WO prophage that is widespread in *Wolbachia* populations^40^. We searched for the presence of prophages in the Ot genomes using the online search tool PHASTER^41, 42^ and identified remnants of prophage genetic material in all strains except for Boryong (**Fig. 5D**, **E**; **Supplementary Dataset 7**). By contrast, no prophage regions were identified in *Rickettsia conorii, Rickettsia rickettsia, Rickettsia prowazekii, Anaplasma phagocytophilum* or *Anaplasma marginale,* although two sites were detected in both the genome of *Wolbachia* endosymbiont of *Drosophila melanogaster* and *Rickettsia bellii.* This suggests that prophages are not universally circulating in Rickettsiales populations, but are present in selected species such as Ot, wolbachiae and *R. bellii.* Whilst many of the Ot prophage genes identified by PHASTER include transposase and integrase genes, which may be of ICE origin rather than phage origin, phage-specific genes including capsid and envelope proteins were also found. In addition to the identification of potential prophage regions found by PHASTER, isolated phage-related genes, such as the phage portal protein previously annotated as a hypothetical protein (Fig. 3F), are also present in the Ot genomes. Sequence similarity searches indicated low similarity to a range of diverse phage sequences from free-living bacteria indicating that either the prophages came from numerous sources, or that the sequences within each strain were sufficiently degraded so they have lost identifiable homology to one another.

### RAGE mobilisation genes

#### F-T4SS

The RAGE encodes a conjugative T4SS highly similar to the F-T4SS of the archetypal F plasmid of *E. coli* (*tra/trb)*^8, 9^. Previous comparisons of the RAGE T4SS with that of the *E. coli* F plasmid showed that it encodes 14 proteins predicted to form the T4SS scaffold, some of which are analogous to components within P-type T4SSs^18, 43^ (**Fig. 6A, B, Supplementary Dataset 8**). While syntenic to the *E*. *coli tra*/*trb* T4SS, the RAGE T4SS lacks genes involved in the regulation of conjugation, as well as other assembly factors and lytic transglycosylases (**Fig. 6C**). In this way, the RAGE T4SS is a streamlined version of the canonical F-T4SS. The Ot RAGE T4SS is also highly similar in gene order and composition to F-T4SSs characterized in the RAGEs of *R. buchneri*^11^*, R. bellii*^12^*, R. felis*^15^ and *R. massiliae*^13^. As with these prior reports, we also did not identify a gene encoding a pilin protein (typically TraA in F-T4SSs) in RAGE T4SSs, though it may be that a pilus is synthesized using a different pilin gene since the RAGE-harbouring *R*. *bellii* forms large pili during host infection^12^. Experiments are needed to determine if the RAGE T4SS elaborates a pilus or functions pilus-less, as is noted for the P-T4SS of *Rickettsia* species^43^, *Neorickettsia risticii*^44^, and likely all Rickettsiales^45^. Another common peculiarity of these F-T4SSs is the split gene encoding TraK, the significance of which is unknown.

**Figure 6.**
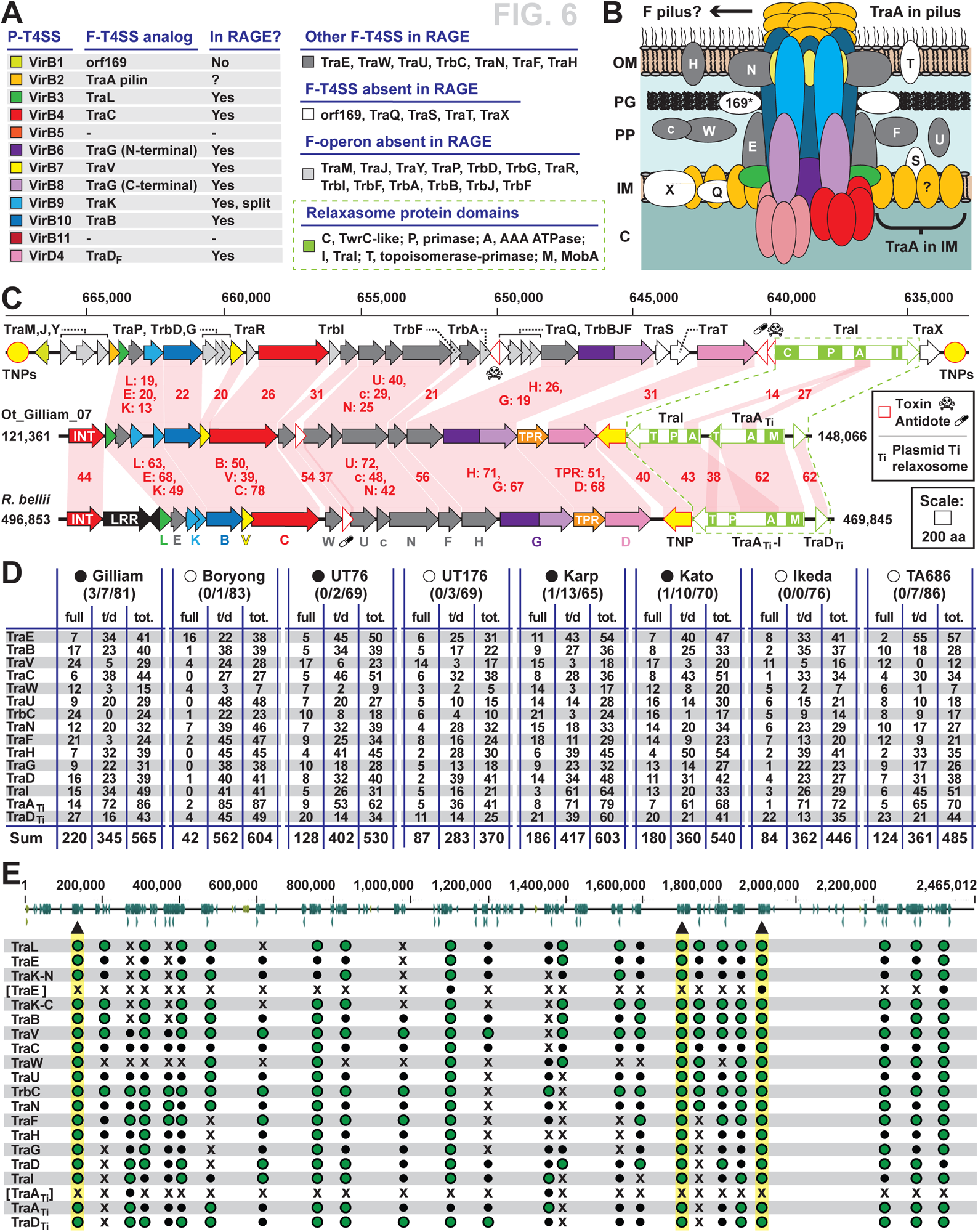
Characteristics of the F-type T4SS and relaxosome proteins encoded on Ot RAGEs. **A**. Composition of the Ot RAGE F-T4SS in relation to the *Agrobacterium tumefaciens vir* P-T4SS and the *Escherichia coli tra/trb* F-T4SS from the F operon. Analogues across divergent T4SSs are coloured similarly, with other colours as follows: dark gray, RAGE T4SS proteins found in F-T4SSs but not P-T4SSs; white, *E*. *coli* F-T4SS scaffold genes not present in RAGE T4SSs; light gray, other *E*. *coli* F operon genes not present in RAGE. For relaxosome proteins (olive green), domains were predicted with SMART ^82^. **B**. Theoretical assembly of the RAGE T4SS in relation to data from other F- and P-type T4SSs. The uncertain synthesis of a pilus is depicted (see text for details). **C**. Comparison of the *E. coli* F operon to mobilisation genes of complete RAGEs from Ot str. Gilliam and *Rickettsia bellii* str. RML369-C. This *E*. *coli* strain, K-12 ER3466 (CP010442), has the F operon on a chromosomal segment flanked by transposases (yellow circles). Red shading and numbers indicate % aa identity across pairwise alignments. Dashed lines enclose the relaxosome genes, whose protein domains are described in panel A. INT, integrase; LRR, leucine rich repeat protein. **D**. Frequency and distribution of of full length and truncated *tra*/*trb* genes in Ot strains. Complete circles, genomes containing full sets of *tra*/*trb* genes within one or more RAGE; open circles, no complete *tra*/*trb* gene sets. Numbers in parentheses: number of complete RAGEs/number of complete RAGE genes containing truncated genes/incomplete RAGEs. Details of truncated genes and gene fusions are given in Supplementary Datasets 1 and 8. **E**. Genomic location of *tra*/*trb* gene clusters in Ot str. Gilliam. Triangles and highlighting depict complete RAGEs. Bracketed TraE and TraA_TI_ are commonly occurring pseudogenized duplications. Green circles, complete gene; small black circles, predicted pseudogene; Xs, gene absent with *tra*/*trb* gene cluster.

#### Relaxosome

The relaxosome of the *E*. *coli tra*/*trb* F-T4SS encodes one multifunctional relaxase, TraI, which excises and binds single-stranded plasmid DNA^46^. In contrast, the Ot RAGE carries three genes *traI*, *traA_Ti_* and *traD_Ti_* predicted to comprise the relaxosome that mobilises RAGE (**Fig. 6C**). *E*. *coli* TraI harbours four distinct domains required for nicking, binding, and unwinding DNA. By contrast, Ot TraI lacks a domain for nicking DNA and shares very limited similarity to *E*. *coli* TraI. However, Ot TraA_Ti_ carries a MobA-like domain that cleaves single- and double-stranded DNA at specific sites^47^. Curiously, all of the domains encompassed by Ot TraI and TraA_Ti_ proteins are found in a single *Rickettsia* RAGE protein, named TraA_Ti_-I, which is highly similar to both Ot TraI and TraA_Ti_ but shares limited similarity to *E*. *coli* TraI. As their annotation indicates, RAGE TraA_Ti_ and TraD_Ti_ are similar to relaxosome proteins of plasmid Ti of *Agrobacterium tumefaciens*, TraA and TraD, which are required for T-DNA translocation into plant cells via the *vir* T4SS^48^. The significance of different relaxosome structures between Ot and *Rickettsia* RAGEs is unclear, although *traA_Ti_* and *traD_Ti_* genes are common on *Rickettsia* plasmids even when RAGE are absent^49^. It may be multiple RAGE types exist in the rickettsial mobilome and are defined by their cognate relaxosomes. The presence of transposases flanking relaxosome genes in all complete Ot and *Rickettsia* RAGEs may also signify that RAGEs evolve by recombining different relaxosome cassettes into the conjugation and cargo genes.

#### Proliferation of Ot RAGE mobilisation genes

Aside from shared synteny and mobilisation gene composition, *Rickettsia* and Ot RAGEs have common insertion points for antidote genes of toxin-antidote modules and transposases (**Fig. 6C**). However, certain *Rickettsia* RAGEs have cargo genes inserted at different sites within the mobilisation genes^11, 15, 18^. Furthermore, a recent study annotated a RAGE from the *Tisiphia* endosymbiont of *Cimex lectularius* that harbours unique mobilisation genes and cargo gene insertion sites^50^. This indicates that RAGEs are far more diverse and widespread across Rickettsiales than previously appreciated. Still, most *Rickettsia* genomes either lack RAGE entirely or show minimal evidence for RAGE insertion near a common genomic position, tRNA^Val-GAC 11^. This is in stark contrast to the proliferated nature of RAGEs in Ot genomes.

The scattershot distribution of RAGE in Ot genomes is particularly evinced by the mobilisation gene clusters that are present in numerous copies within the plethora of RAGEs. We find that over 50% of the RAGE T4SS and relaxosome genes are present as truncated pseudogenes, and that some of these clusters encode only a subset of the 18 possible Ot RAGE mobilisation genes (**Fig. 6D, E**). Given the high degree of pseudogenization, we sought to examine whether any strain encoded any RAGE mobilisation gene clusters containing a complete complement of full-length genes. We found that Karp, Kato, Gilliam and UT76 encoded at least one complete RAGE mobilisation gene set, whilst Ikeda, Boryong, TA686 and UT176 did not (**Fig. 6D, E**, **Supp. Fig. 3, Dataset 7**). There was a positive correlation between strains containing complete sets of RAGE mobilisation gene sets and the total number of full-length mobilisation genes (**Fig. 6D, E**). Moreover, the complete RAGE mobilisation gene clusters in Gilliam and Kato were located within complete RAGE regions (**Fig. 1C**, **Fig. 6E**, **Supp. Fig. 3**). Whilst several genomes lack complete RAGE mobilisation gene clusters, all except Boryong encode at least one full length copy of each RAGE mobilisation gene, albeit not in a contiguous cluster. Therefore, it is possible that all strains except Boryong could assemble a functional F-type T4SS competent to mediate transfer of RAGEs.

### The impact of pervasive mobile genetic elements on the Ot genome

#### P-T4SS

Like other rickettsial species, Ot encodes a P-type T4SS related to the archetypal *vir* T4SS of the pTi plasmid of *A. tumefacians*. Relative to *vir*, this **Rickettsiales *vir* homolog** *(**rvh**)* T4SS has distinct features, including the scattered distribution of *rvh* gene clusters, duplication of *rvhB8*, *rvhB9* and *rvhB4,* 3-5 copies of *rvhB6*, and no gene encoding an equivalent to the VirB5 minor pilin subunit^45, 51, 52^(**Fig. 7A,B**). These characteristics are nuanced: 1) RvhB4-II, RvhB8-II, and RvhB9-II carry atypical structural deviations from described VirB4, VirB8, and VirB9 family proteins, 2) RvhB6 proteins have large insertions flanking the VirB6-like membrane spanning region, and 3) a lack of a minor pilin subunit precludes formation of a T-pilus^52, 53^. There is evidence that structurally different RvhB8-I and RvhB8-II proteins of *R*. typhi cannot dimerize^53^, which led to the hypothesis that divergent duplications may autoregulate effector secretion^52^. The recent identification of *rvh* genes in all seven Rickettsiales families implies a highly important function^3, 54^. Thus, we assessed the properties of the Ot *rvh* T4SS in the face of its rampant mobile genetic element-induced genome shuffling.

**Figure 7.**
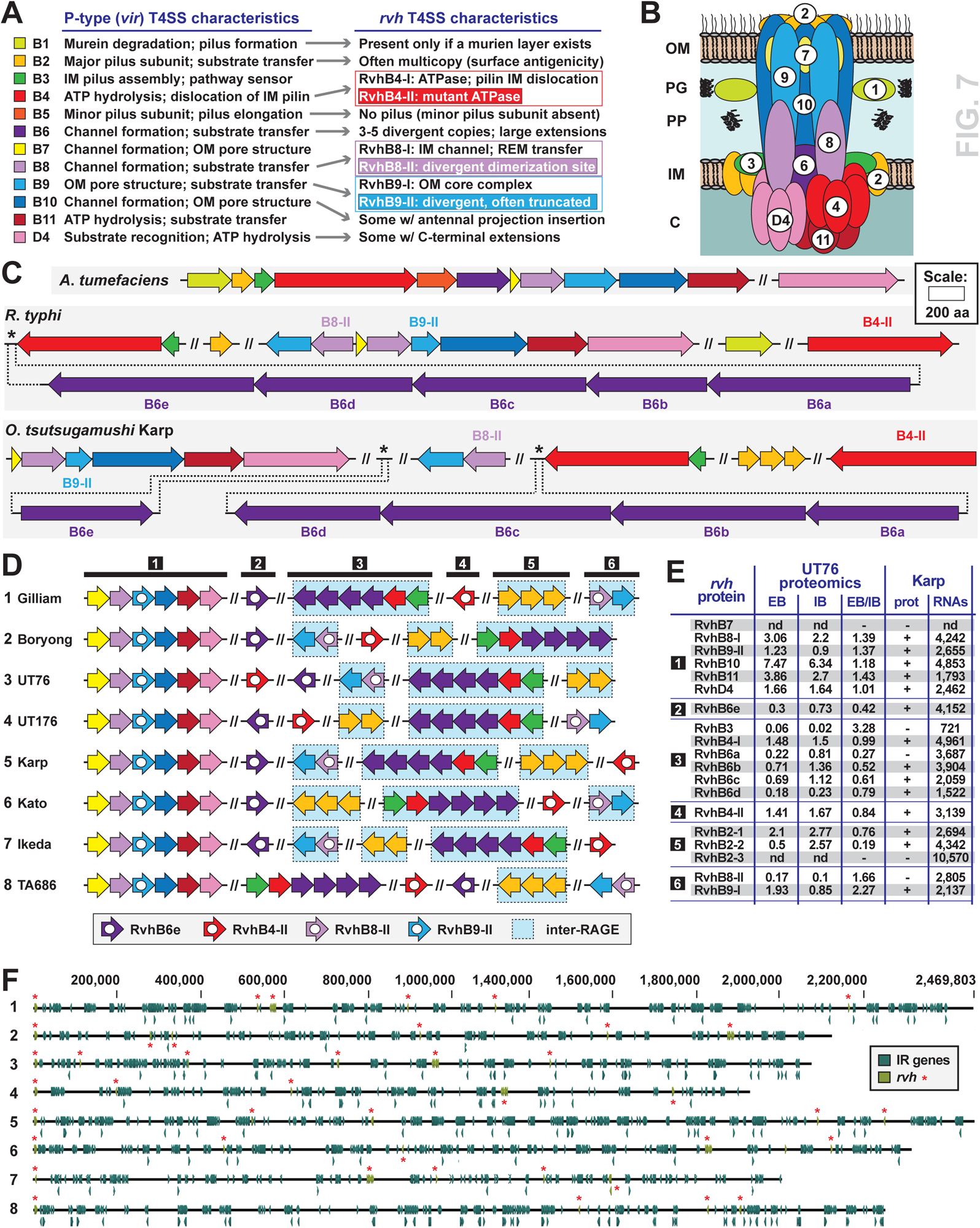
Synopsis of Ot P-type (*vir*-like) T4SS genes. **A.** Description of the general *rvh* T4SS characteristics, summarized from prior studies^43, 45, 51, 52^. **B**. Theoretical assembly of the RAGE T4SS in relation to data from other P-type T4SSs. There is no synthesis of a T-pilus (see text for details) **C**. Comparison of genes encoding *vir* T4SS in *Agrobacterium tumefaciens,* the archetypal P-T4SS, and those encoding the *rvh* T4SS in *Rickettsia typhi* and Ot. **D**. Arrangement of *rvh* genes in Ot genomes. Red genes are located in RAGE regions whilst blue are located in IR regions. **E**. Previously published RNAseq and proteomics data showing relative expression levels of *rvh* genes in strains UT76 and Karp. These are taken from Atwal et. al 2022 (UT76) and Mika-Gospodorz et. al 2020 (Karp). UT76 data shows relative peptide counts in intracellular bacteria (IB) and extracellular bacteria (EB). Karp data shows presence or absence of detectable peptides from proteomics analysis (+/-) and relative RNA transcripts from RNAseq data (TPM/transcripts per million). **F**. Distribution of *vir* genes across Ot genomes showing lack of conservation of absolute position, despite similarities in gene groupings as shown in **Fig. 7D**.

A single set of *rvh* genes is present in all the Ot genomes analysed here and has features resembling those described in other Rickettsiales species (**Fig. 6A-C**). Interestingly, Ot shares two key characteristics with the *rvh* T4SS of distantly-related Anaplasmataceae species as opposed to that of closely-related *Rickettsia* species. First, while Ot lacks genes for a VirB5 protein and therefore cannot form extracellular pili used for cellular attachment, it encodes 2-3 copies of *rvhB2*, the major pilus subunit. Given that Ot lacks **lipopolysaccharide** (**LPS**) on its surface, multiple RvhB2 proteins may act as divergent surface antigens in a similar fashion previously posited for Anaplasmataceae species, which collectively lack LPS biosynthesis genes and have multiple *rvhB2* genes throughout their genomes^45, 55^. Second, Ot lacks any identifiable *rvhB1* gene, which is present only in *Rickettsia* spp. This gene encodes a lytic transglycosylase predicted to cleave **peptidoglycan** (**PGN**) to allow T4SS scaffold assembly^51^. Compared with *Rickettsia* spp., which synthesize a canonical PGN layer^56, 57^, the presence of a minimal cell wall in Ot and Anaplasmataceae species is consistent with the absence of *rvhB1* from these genomes^58, 59^. These collective differences in *rvh* T4SS architecture present clear convergent evolution in Ot and Anaplasmataceae species in the context of shared cell wall morphology and probable responses to host cell immune pressures^60^.

Our analysis shows that the identities of six *rvh* gene clusters are conserved across Ot strains, whilst the genomic positions of the clusters vary between strains (**Fig. 7D**). Clusters 1 (*rvhB7*, *rvhB8-I*, *rvhB9-II*, *rvhB10*, *rvhB11*, *rvhD4*), 2 (*rvhB6e*) and 4 (*rvhB4-II*) are located within RAGEs in all strains, with the other *rvh* genes consistently located in IR regions except for cluster 6 and clusters 3 and 6 in UT176 and TA686, respectively (**Fig. 7D, F**). Analysis of published datasets of proteomics and RNAseq data in Karp and UT76^19, 22^ show that RvhB2-3 and RvhB7 proteins are not detected under growth conditions used in those analyses, although transcription levels of *rvhB2-3* are high (**Fig. 7E**). All the other Rvh proteins are detected in UT76 and most are detected in Karp. The UT76 dataset compared peptide levels in two different bacteria populations: **intracellular bacteria** (**IB**) and **extracellular bacteria** (**EB**)^22^. The EB/IB ratio of some multi-copy Rvh proteins differs between paralogs; e.g., RvhB2-1, which is present at a ratio of 0.76 compared with the ratio of 0.19 for RvhB2-2. This suggests expression of these proteins may be differentially regulated, potentially reflecting functional differences. Our collective analyses indicate that Ot Rvh genes can form a functional P-T4SS, despite the pervasive mobile element-induced gene shuffling in Ot genomes. While no Ot *rvh* transported effector has been described to date, it is highly likely that Ot utilizes the *rvh* T4SS during host cell infection, as secreted proteins that interact with the T4SS gatekeeper, RvhD4, have been described for *R. typhi*^61–63^*, R. rickettsii*^62, 64^*, A. marginale*^65^*, A. phagocytophilum*^66–69 70–72^ and *Ehrlichia* chaffeensis^68, 73 74–76^.

### Ot lacks defence mechanisms against invasive DNA

We show here that the Ot genome is exceptional in its abundance of invasive mobile genetic elements, including ICEs, transposases, group II introns and prophages. Bacteria have evolved a range of anti-viral mechanisms to minimise damage caused by mobile genetic elements^77–79^. We therefore sought to assess if Ot lacks these protective systems, possibly explaining the proliferation of mobile genetic elements. We used DefenseFinder to carry out a systematic search for all known anti-phage systems including restriction modification systems, CRISPr/Cas systems, and toxin-antidote defence modules^78, 80^. We found that none of the Ot strains in our study had any identifiable defence systems. Whilst it is possible that this is due to sequence divergence, small size (e.g., certain toxin-antidote modules) or systems that have not yet been discovered, the software was able to detect three different restriction modification systems and the newly described Pyscar defence system^81^ in the closely related free living alphaproteobacterial *Caulobacter crescentus.* In addition to lacking identifiable antiviral defence systems, Ot also has limited homologous recombination capability, a system that is frequently used in antiviral defence ^78^. Whilst Ot encodes RecA and the alternative homologous recombination pathway RecFOR, it lacks the major repair complex RecBCD that can defend against some mobile genetic elements by degrading linear double stranded DNA^10^. Overall, Ot lacks identifiable mobile genetic elements defence systems likely explaining the proliferation of mobile DNA in these genomes.

## Conclusions

The identification of complete RAGEs in two Ot strains raises the possibility that these ICEs are active at the population level. Evidence for this hypothesis awaits whole genome sequencing of large numbers of Ot isolates beyond the total of eight currently available. Whilst only two genomes encode complete RAGEs with full length genes, all encode all the genes required for RAGE mobilisation, albeit in dispersed locations across the genome. Future research is needed to determine whether such mosaic RAGEs can be mobilised or not.

The identification of potentially active RAGEs in Ot raises the question of how they can be transferred between Ot organisms during their lifecycle. Ot is an obligate intracellular bacterium and therefore bacterial cells have limited interactions with other bacteria of the same or different species. It is possible that different strains of Ot infect the same host cell in a mite or a rodent during co-infection by two species. Albeit rare, this could occur with sufficient frequency to enable horizontal gene transfer between species. Alternatively, it is possible that the extracellular form of Ot retains sufficient residual metabolic activity to support lateral DNA transfer in the cell-free extracellular state. Mites typically feed in a tight cluster, for example on the ear of an infested rodent, and the co-feeding pool may provide the environment for close encounters between Ot cells in an intracellular or extracellular state to mediate conjugation. Finally, while not detected in other environments or hosts (i.e. protists), there could be other opportunities for Ot strains to exchange DNA or acquire DNA from other intracellular species.

In conclusion, this study has led to the manual re-annotation of the genomes of eight strains of Ot, enabling the delineation of RAGE and IR regions. Open questions remain. Importantly, whilst intact RAGEs have been identified in two strains, the dynamics of the Ot RAGE are completely unknown. It is also unknown whether the current set of RAGEs within one genome results from one or multiple invasion events. The Ot RAGE encodes an F-T4SS, but it is not known if these are active, nor what they transport beyond the ICE itself. Progress towards answering these questions will enable further insights into the biology and pathogenicity of this important human pathogen.

## Methods

**Table Methods 1:** Accession numbers of genomes used in this study

| Strains | Genome accession numbers | Links |
| --- | --- | --- |
| **Boryong** | AM494475.1 | https://www.ncbi.nlm.nih.gov/nuccore/AM494475.1 |
|  | NC009488.1 | https://www.ncbi.nlm.nih.gov/nuccore/NC_009488.1 |
| **UT76** | LS398552.1 | https://www.ncbi.nlm.nih.gov/nuccore/LS398552 |
| **UT176** | LS398547.1 | https://www.ncbi.nlm.nih.gov/nuccore/LS398547.1 |
| **Karp** | LS398548.1 | https://www.ncbi.nlm.nih.gov/nuccore/LS398548.1 |
| **Kato** | LS398550.1 | https://www.ncbi.nlm.nih.gov/nuccore/LS398550.1 |
| **Ikeda** | AP008981.1 | https://www.ncbi.nlm.nih.gov/nuccore/AP008981.1 |
|  | NC_010793.1 | https://www.ncbi.nlm.nih.gov/nuccore/NC_010793.1 |
| **TA686** | LS398549.1 | https://www.ncbi.nlm.nih.gov/nuccore/LS398549.1 |
| **Gilliam** | LS398551.1 | https://www.ncbi.nlm.nih.gov/nuccore/LS398551.1 |

### Identification of RAGE and inter-RAGE regions in the genome of Ot

The boundaries of RAGE and IRs were manually delineated in each genome using defined criteria. First, groups of genes whose relative position to one another was conserved across strains were identified manually by comparing the genomes of 8 Ot strains (Ikeda, Boryong, Karp, Kato, Gilliam, TA686, UT76 and UT176). This led to the identification and numbering of IR regions. RAGE regions were subsequently identified using criteria largely drawn from *K Nakayama et al*, 2008^9^.

The element was classified as a “complete RAGE gene” if the sequences encoded a full-length integrase gene at the left end (N-terminus), a full-length transposase gene, full-length set of conjugative transfer genes (tra genes: *TraA, TraB, TraC, TraD, TraE, TraF, TraG, TraH, TraI, TraK, TraL, TraN, TraU, TraV, TraW*), and nonconjugative genes (RAGE associated cargo genes) including one or all of the following: SpoT-related proteins (ppGpp hydrolase, (p)ppGpp synthetase, SpoT synthase, and SpoT hydrolase), DNA methyltransferase, DNA helicase, histidine kinases, ATP-binding proteins (mrp), HNH endonuclease, membrane proteins, ankyrin repeat proteins, and hypothetical proteins. The RAGE associated cargo genes in “complete RAGE gene” can be either full-length or truncated genes.

The element was classified as an “complete RAGE with truncated genes” if the sequence encoded the same gene set as above, but where one or more of the integrase, transposase, or Tra conjugative transfer genes were truncated.

The element was classified as an “incomplete RAGE” if the sequence encoded integrase or transposases, and at least one RAGE associated cargo gene.

The “isolated mobile gene” was defined as encoding one or more integrase or transposases without RAGE associated cargo genes.

The “isolated cargo gene” was defined as encoding one or more cargo genes without transposases, integrases, or Tra genes.

The “isolated hypothetical protein” was defined as encoding one or more hypothetical proteins at the boundary of conserved IRs or RAGEs.

The presence of a *dnaA* gene was used as an indicator gene for defining the end of a RAGE element (Fig. Methods 1). However, the criteria could not be applied for all RAGE elements when hypothetical proteins and transposases are located at the end of RAGE masking the original *dnaA* terminus. In the first case (Fig. Methods 1A), RAGE elements are located next to each other in the same direction. RAGE is terminated when integrase gene of the next RAGE is found. In the second case (Fig. Methods B), RAGE elements are located next to each other in opposite direction and two *dnaA* genes are located next to the each other. In this case the RAGE is terminated at the *dnaA* gene which belongs to RAGE on the left (forward direction) and RAGE on the right (reverse direction). In the third case (Fig. Methods C), RAGE elements are located next to each other in opposite direction. Two RAGEs were combined into one RAGE if a *dnaA* gene was not present in either RAGE.

**Figure Methods 1.**
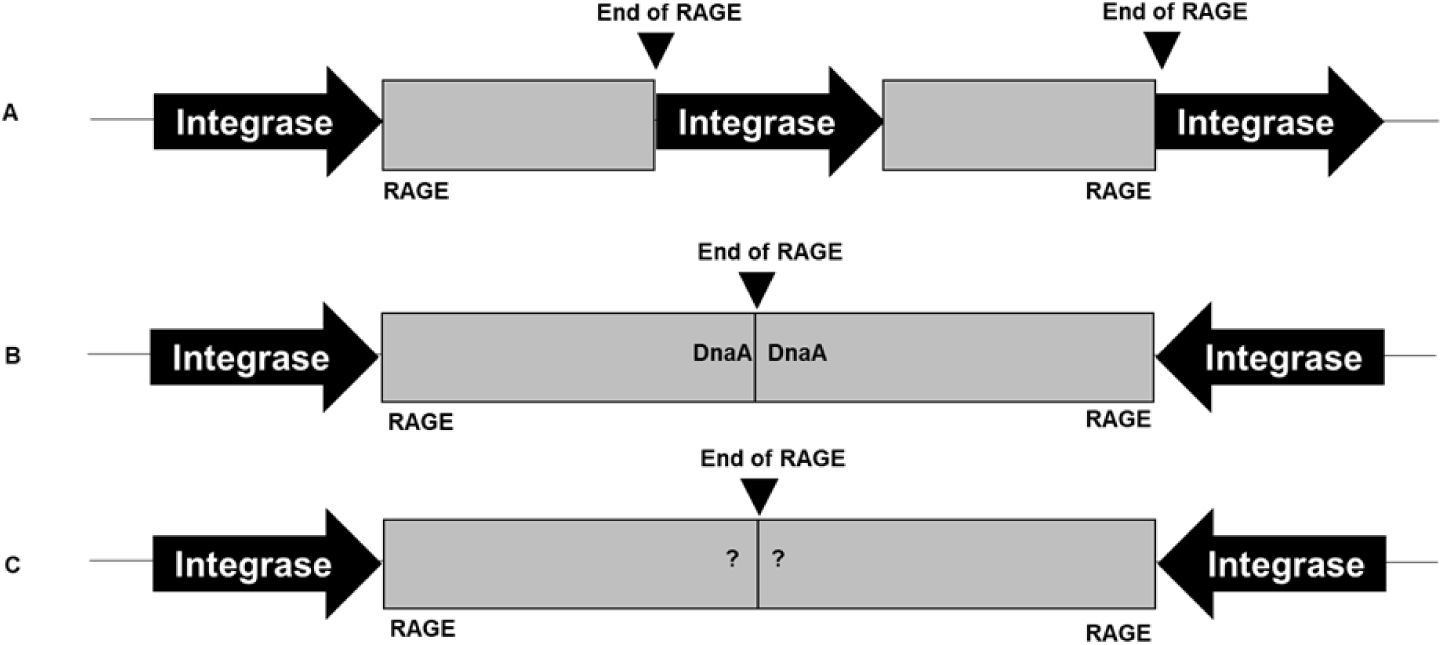
Identification of RAGE termination positions.

### Identification of RAGE associated cargo genes

The list of conserved nonconjugative genes or RAGE associated cargo genes were extracted based on *K Nakayama et al*, 2008^9^. The sequences were then inspected manually by observing the length of genes, conserved motifs/domains, and other elements such as signal peptides, transmembrane regions etc. A gene was defined as a “Full-length gene” if the sequence encoded a complete set of domains, a “Truncated gene” if the sequence encoded only a partial set of domains and a “Degraded gene” if no domain was identified on the sequence.

### Analysis of multicopy cargo genes not associated with DNA mobilisation

#### Membrane proteins

Membrane proteins were manually extracted from the genome database and the SMART search engine^82^ was used to identify membrane domains and other elements. The membrane protein was assigned as “OT_RAGE_membrane_protein” if no identifiable domain was identified. The membrane protein was assigned on new name if significant domain/main domains were found such as autotransporter proteins (Sca family), vut1-Putative vitamin uptake transporter, RhaT-Permease of the drug/metabolite transporter (DMT) superfamily, and Bax inhibitor-1 (BI-1)/YccA inhibitor of FtsH protease domains.

#### MRP/histidine kinases

All genes previously annotated as histidine kinase (HK) and multidrug resistance-associated proteins (MRPs) were extracted from the genome databases and analysed using SMART^82^. Both HK and mrp proteins contain an HATPase_c domain (Histidine kinase-type ATPases catalytic domain). This domain is found in several ATP-binding proteins, including: histidine kinase^83^, DNA gyrase B^84^, topoisomerases, and heat shock protein HSP90^85^. The new naming system of HK and mrp were classified based on the HATPase_c domain. HK or mrp proteins were renamed as “HATPase_c domain containing protein” if the search found an HATPase_c domain. HK or mrp proteins were renamed as “degraded HATpase_c” if no significant domains were found on the search. HK and mrp proteins in this study were classified as truncated gene because they only contained catalytic part (HATPase_c) and lacked other major domains such as sensor domain, HisKA (Histidine kinase A domain dimierization/His phospotransfer), and receiver Hpt (Histidine phosphotransfer)^86^.

**Fig. Methods 2.**
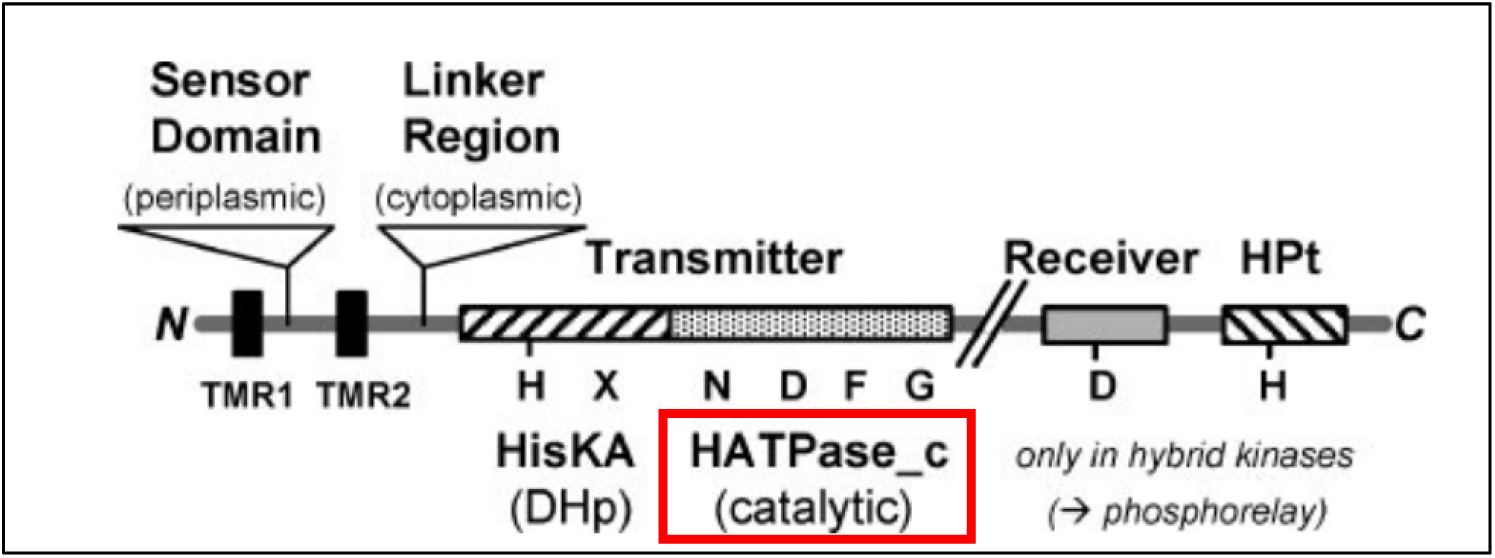
The overview of domain of histidine protein kinases. Figure is modified from ref^86^.

In addition, the same HATPase_c domain was also found in symporter proteins. The sodium:proline symporter “PutP” was classified as full-length if the sequence contained symporter, HisKA, HATPase_c, and REC domains. PutP was was classified as “truncated PutP” or “degraded PutP” if the sequence lacked the symporter and REC domains. The symporter Sodium:pantothenate symporter “PanF” was classified as full-length if the sequence contained only the symporter region.

#### (p)ppGpp hydroplase/synthetases

We manually extracted all genes annotated as: SpoT, (p)pGpp, synthetase, and hydrolase from the genome databases. The sequences were then compared to their respective orthologs in *Escherichia coli* (ECO) and *Caulobacter crescentus* (CCS). Literature searches and GenomeNet motif search (pfam) were used to identify motifs and conserved regions in these sequences as shown (**Table Methods 2).**

**Table Methods 2.**
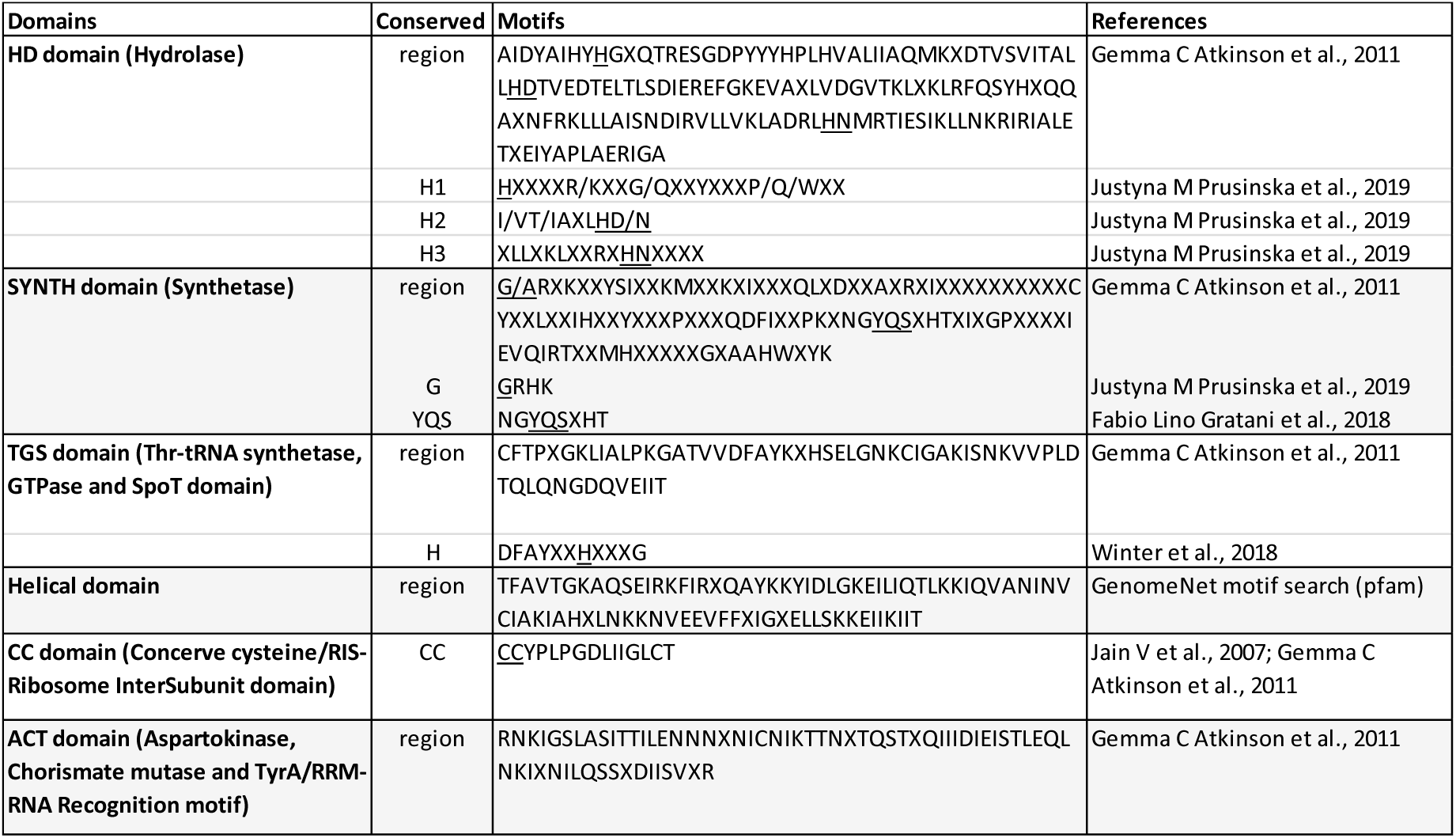
Motifs conserved in SpoT. <u>Underlined</u> base(s) indicate important amino acid in the motif.

We employed a naming system based on the presence of domains/motifs in each CDS. genes containing all domains (HD, SYNTH, TGS, Helical, CC, ACT) were annotated as SpoT. Genes having only the HD domain were annotated as SpoT-hydrolase, genes encoding only SYNTH domain was annotated as SpoT-synthetase. Genes encoding the HD domain but lacking one or more of the conserved histidines was annotated as truncated hydrolase. Short fragments that could be aligned to Hydrolase but lack complete domains were annotated as degraded hydrolase. Genes containing HD domain merging with a part of HATPase were named as HATPase-SpoT-hydrolase and genes containing HD domain merging with a part of Mrp were named as Mrp-Hydrolase, respectively.

#### DNA methyltransferases

DNA methyltransferase (MTase) genes were extracted from the genome and analysed on SMART^82^ for protein domain annotation. However, SMART does not provide the details of MTase motifs within the predicted domain. Therefore, multiple sequence alignment of DNA methyltransferase was further characterized for identification motifs using Geneious. We used the conserved amino acid residues in the Dam (DNA adenine methyltransferase) protein of *E. coli* (acc.no. P0AEE9) as a reference for identification motifs I-VII & motif X of MTase at C-terminal region^87^. The protein sequence of DNA methyltransferase containing motif I-VII & motif X was indicated as full-length gene. The protein sequence of DNA methyltransferase with incomplete motifs and unidentified motifs were indicated as truncated gene, and degraded gene, respectively.

#### Replicative DNA helicases

DNA helicase genes were filtered from the genome and their protein domains were characterized by SMART^82^. Multiple sequence alignments of the helicase genes was then carried out in order to identify motifs. We used the conserved amino acid residues in the DnaB protein of *E. coli* K12 (acc.no. NC000913.3) as a reference for the identification of motifs I-VII at C-terminal region ^88^. The protein sequence of DnaB containing motif I-VII was indicated as full-length gene. The protein sequence of DnaB with incomplete motif and unidentified motifs was indicated as truncated gene and degraded gene, respectively.

#### Uncharacterised proteins

Between 308-464 genes annotated as hypothetical or uncharacterised were found in the eight genomes of *Orientia*. In this study, we only manually analysed uncharacterised proteins from Karp strain as a model to minimize the analysis time. Uncharacterised proteins from Karp were filtered from the genome and the protein domain was characterized by SMART^82^. Where clear groups of homologous genes within the set of Karp genes was found, these were classified into 25 defined groups. These were renamed according to known domains with which they had homology, or named Ot_RAGE_hypo_group 1-7. These 25 groups were then aligned to the other seven genomes in our dataset in order to determine the conservation of the groups of genes.

Some uncharacterised proteins were changed to new name, and no longer classified as hypothetical proteins, if they aligned to known genes such as DnaA, Phage portal protein, Lipase3 etc.

### Analysis of multicopy genes involved in DNA mobilisation

#### Insertion sequence transposable elements

The presence of insertion sequence (IS) elements in Ot was investigated using the online search tool ISfinder^89^ to match with attributes and nomenclatures previously submitted for Orientia-specific IS^9^. Each IS match was manually traced for completeness with flanking inverted repeats (IR) and direct repeats (DR) along respective genome sequences. Extensive analysis was performed with Karp strain to identify the complete set of IS elements and classified into classes. Only ISOt3 and ISOt5 were systematically analysed across all 8 different *Orientia* genomes.

#### Integrases

Genes annotated as integrase genes were extracted from the genomes and protein domains were screened by SMART^82^. Integrase in *Orientia* is a phage integrase which is classified into two major families: the tyrosine recombinases and the serine recombinases, based on mode of catalysis^90^. Then multiple sequence alignment of phage integrase domain was analysed for identification motifs using Geneious. We used the conserved amino acid residues in Bacteriophage P2-integrase (acc.no. AF063097.1) and Enterobacteria phage P2-integrase (acc.no. NC_009488.1) as references for the identification of three domains; arm-type binding motifs at N-terminal region, core-type binding (CB), and catalysis at C-terminal region. The His-X-X-Arg motifs and second conserved Arginine on catalytic domain were also included in the alignment^90, 91^. The protein sequence of phage integrase containing arm-type binding, core-type binding, and catalysis motifs was indicated as full-length gene. The protein sequence of phage integrase with incomplete motif and unidentified motif was annotated as truncated gene and degraded gene, respectively.

#### Reverse transcriptases

Reverse transcriptase (*rvt*) genes were filtered from the genome and protein domains were characterized by SMART^82^. Multiple sequence alignment of *rvt* was then analysed for identification motifs. We used the conserved amino acid residues of group II intron reverse transcriptase/maturase (LtrA) in *E. coli* (acc.no. WP_096836589.1), and *Lactococcus lactis* (acc.no. NZ_CP059048.1) as a reference for the identification of three domains; reverse transcriptases (RVT_N) at N-terminal site, reverse transcriptases (RT), and Group II intron, maturase-specific domain (GIIM) at C-terminal site^92–94^. The protein sequence of reverse transcriptase containing RVT_N, RT, and GIIM was indicated as full-length gene. The protein sequence of reverse transcriptase with incomplete motif and unidentified motif was indicated as truncated gene and degraded gene, respectively.

#### Transposases

Genes annotated as transposase genes were first extracted from the genome. Then, protein domains and motifs were further characterized by SMART^82^ and Geneious, respectively. Transposase gene in *Orientia* belong to restriction endonuclease-like proteins or PD-(D/E)XK nucleases and DD[E/D]-transposase, which generally contain the catalytic domain, and transposon-binding domain^95, 96^. Some transposases additionally contain C-terminal or N-terminal domains^97^. In this study, we used the conserved amino acid residues of in *E. coli* (acc.no. NC 002695.2) as a reference for identification restriction endonuclease-like motifs (I-IV). Three conserved active sites in motif II and III, one of which is aspartic acid (D), one is either glutamic (E) or aspartic acid (D) and/or the last one is lysine (K), were identified for characterization of PD-(D/E)XK nucleases and DD[E/D]-transposase motifs ^95, 96^. The protein sequence of transposase containing motifs (I-IV) and PD-(D/E)XK signature residues was indicated as full-length gene. The protein sequence of transposase with incomplete motif and unidentified motif was indicated as truncated gene and degraded gene, respectively.

#### Prophage genes

Potential prophage sequences within 8 *Orientia* genomes were identified using PHASTER (PHAge Search Tool Enhanced Released)^41^ where specific phage related proteins such as ‘coat’, ‘fiber’, ‘head’, ‘plate’, ‘tail’, ‘integrase’, ‘terminase’, ‘transposase’, ‘portal’, ‘protease’ or ‘lysin’ within bacterial genomes were recognized using a sequence identity search.

### Analysis of Ankyrin repeat containing proteins

To identify Ankyrin repeat (AR) proteins (Anks) from previously annotated records, all Ank sequences were extracted and analyzed using SMART^82^ to predict ARs and other domains including coiled-coil, F-box, and PRANC. SMART^82^ defines an Ank as a 33-residue motif. Ank commonly involves in protein-protein interaction. The core of the repeat seems to be a helix-loop-helix structure. SMART’s consensus for an ankyrin repeat is shown in Fig. Methods 3. It is important to note that the protein structure or functionality of any Ank was not characterized in this study. Therefore, any Ank that contains only one or two repeats may be non-functional.

**Fig. Methods 3.**
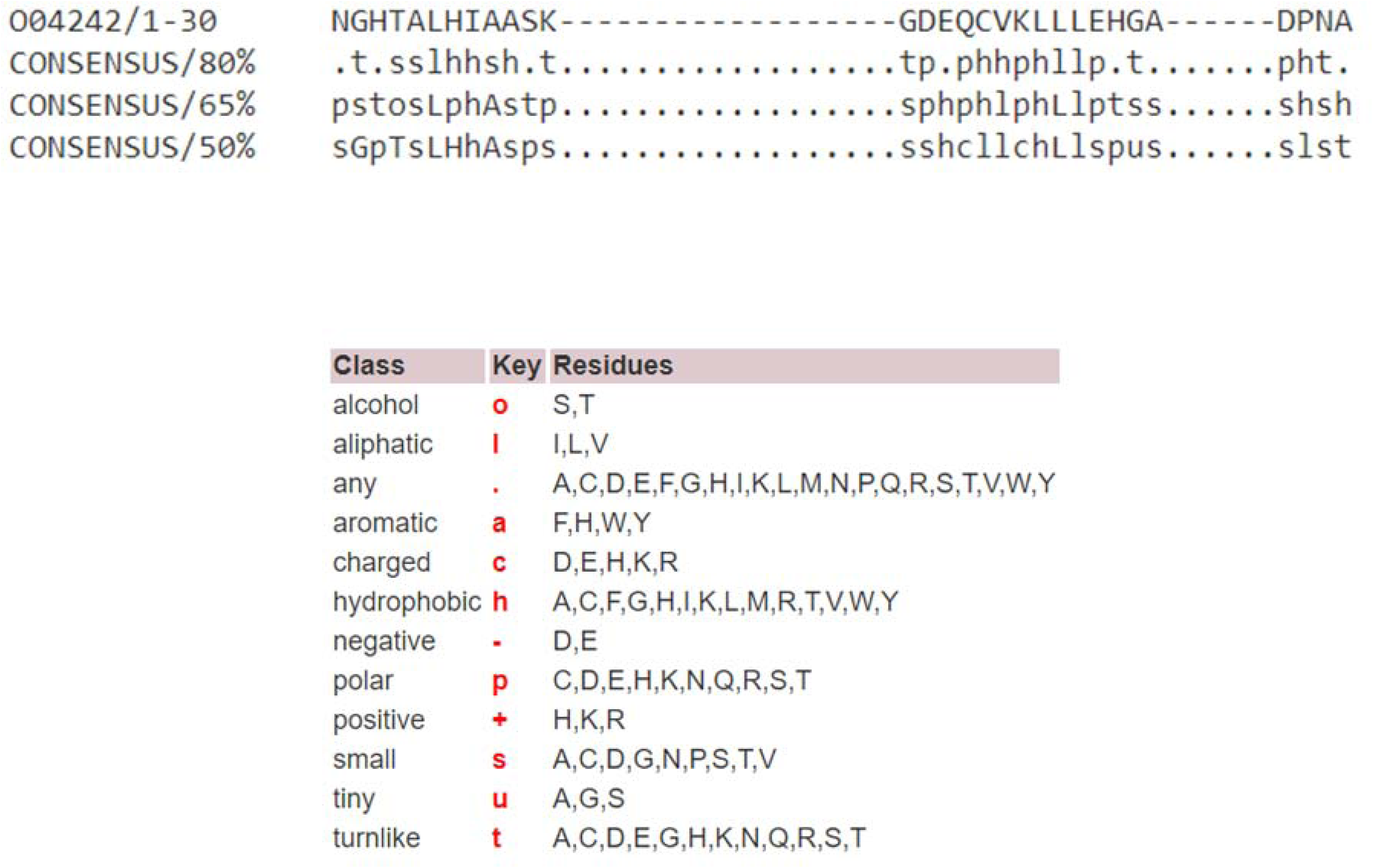
Consensus term for Ank domain by SMART.

Identification of homologous Ank repeats across Ot strains were manually inspected using Geneious. The criteria for identification of a homolog Ank is based on sequence similarity and repeated units. Individual Ank sequences of the other 7 strains were blasted against Ot strain Ikeda. Then, the sequence that presented the highest identity (>80-90%) was chosen to verify the similarity of Ankyrin repeats and other domains. The sequences were given the name based on the published Ikeda Anks if the overall sequence’s identity to Ikeda Anks was more than 80% and presented a similar set of ARs. The sequences were given a new name if the overall sequence identity to Ikeda Anks was less than 80% and presented a different set of ARs, this included extra repeated units or missing repeated units.

Unidentified Anks or hypothetical proteins were manually searched and inspected using Geneious. To search unidentified Anks in Ikeda and other 7 strains, each repeat unit of an individual published Ikeda Anks were imported into “Find motifs” tool, and the maximum mismatches were set up to 10. The closely matched sequences were further identified the repeated units and other domains by SMART. Then, the sequences of unidentified Ank were blasted against the published Anks or blasted within the strain to check whether it was different from identified Anks or not. The newly identified Anks were given a name by continued ranking after the published Ikeda Anks, starting with Ank21, Ank22, Ank23, etc.

### Analysis of Tetratricopeptide repeat containing proteins

To identify TPR proteins from the previous annotated records of Orientia 8 strains, all TPR sequences were extracted and analyzed using SMART^82^. The program predicted the location of TPR motifs and other domains including signal peptide and transmembrane region. All identified TPR proteins were grouped based on the similarity of the location of TPR motifs. Each group consists of one long gene, the master gene, and multiple shorter duplication remnants. For unidentified TPR were manually searched using Geneious. All TPR proteins were renamed based on the group number followed the number of TPR repeats.

### Analysis of P-type IV secretion systems (*rvh)*

Literature search and blast search (NCBI and KEGG) were performed to identify the presence of each Rvh subunit (RvhB1 to VirB11 and RvhD4) in Ot. The amino acid sequences of each Vir subunit present in Ot Boryong strain (OTS) were compared to their respective orthologs in *Rickettsia bellii* (RBE) and *Agrobacterium tumefaciens* (ATU) for their percent of amino acids identity, length of amino acid sequences, and presence of motifs.

The presence of motifs was used as the major criteria to identify Vir subunits as indicated in Table Methods 3:

**Table Methods 3.**
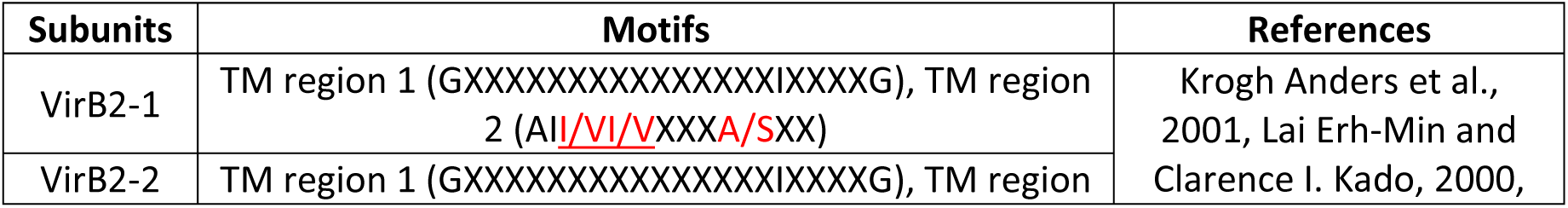
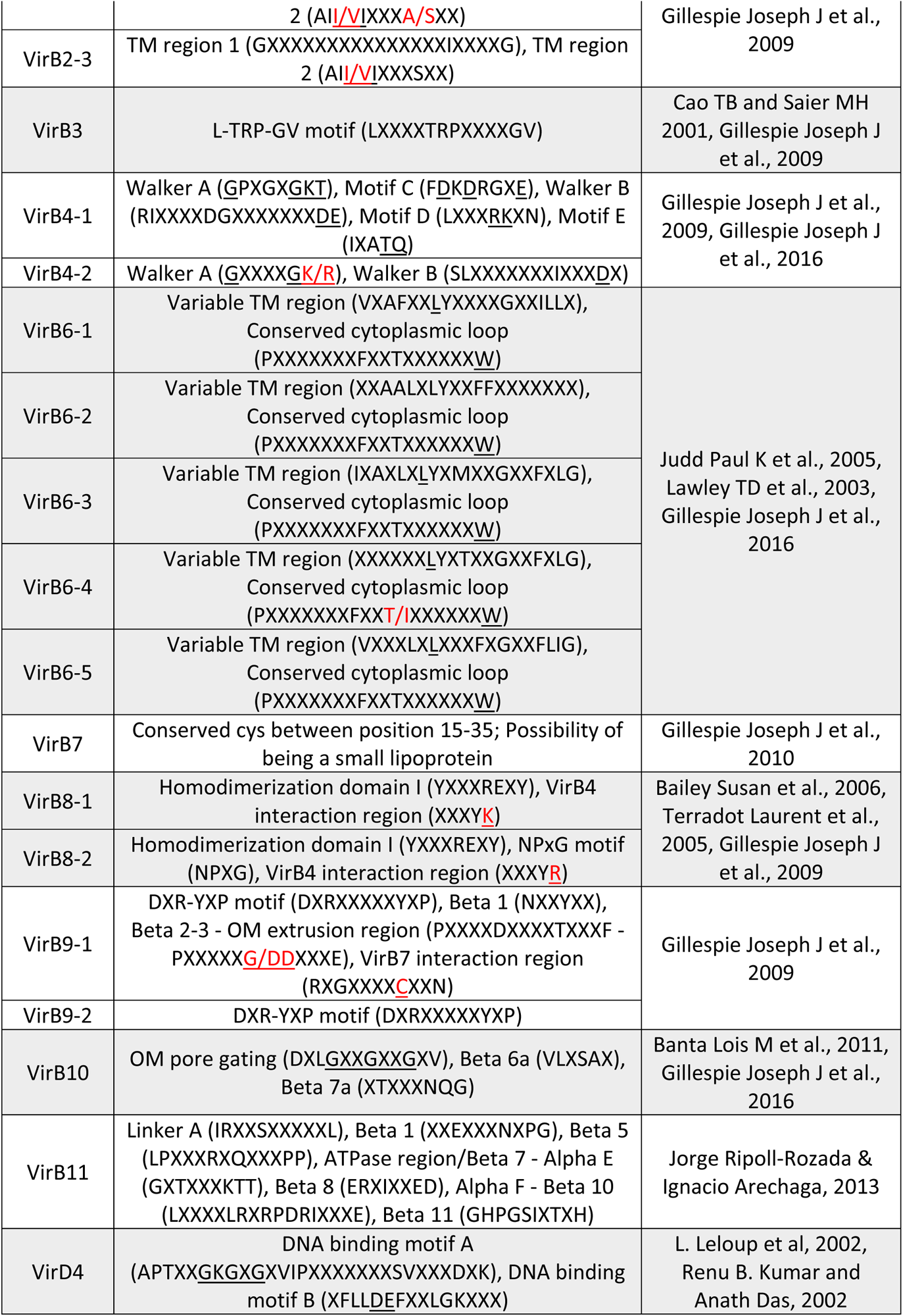
Vir proteins and their motifs. Underlined base(s) indicate important amino acid in the motif. Red letter indicates variability in amino acid sequence.

VirB1 and VirB5 could not be identified in *Ot*. However, VirB7 was annotated based on gene positioning, conserved cysteine(s), and its important role of being a small lipoprotein in T4SS of *Rickettsia* species.

These *rvh* genes identified in Boryong were then set as a reference strain for Ot. Each *vir* gene was then blasted (nucleotide blast using Geneious Prime) to identify the presence of each *rvh* across the eight strains of Ot (Boryong, Ikeda, Karp, Kato, Gilliam, TA686, UT76, UT176). The amino acid sequences of each Rvh subunit from the eight strains of *Ot* were then aligned (Multiple alignment - Clustal Omega) to verify their motifs. Even though some of the gene copies appear to be a truncation or pseudogene due to loss of some motif(s) like that in RvhB4-II, RvhB8-II, and RvhB9-II, they are well characterized in literature. So, these names were kept the same in our annotation.

### Analysis of F-type IV secretion systems (RAGE T4SS)

Literature search and blast search (NCBI and KEGG) were performed to identify the presence of each Tra subunit (F-type T4SS: TraA to TraN, TraU to TraW, TrbC, TrbE and P-type T4SS: TraA_Ti_, TraD_Ti_,) in Ot. The amino acid sequences of each Tra subunit present in Ot Ikeda strain (OTT) were then compared to their respective orthologs in *Rickettsia bellii* (RBE) and *Escherichia coli* (ECZ) for F-type T4SS or *Agrobacterium tumefaciens* (ATU) for P-type T4SS. The presence of motifs was used as the major criteria to identify Tra subunits as indicated in Table Methods 4:

**Table Methods 4.**
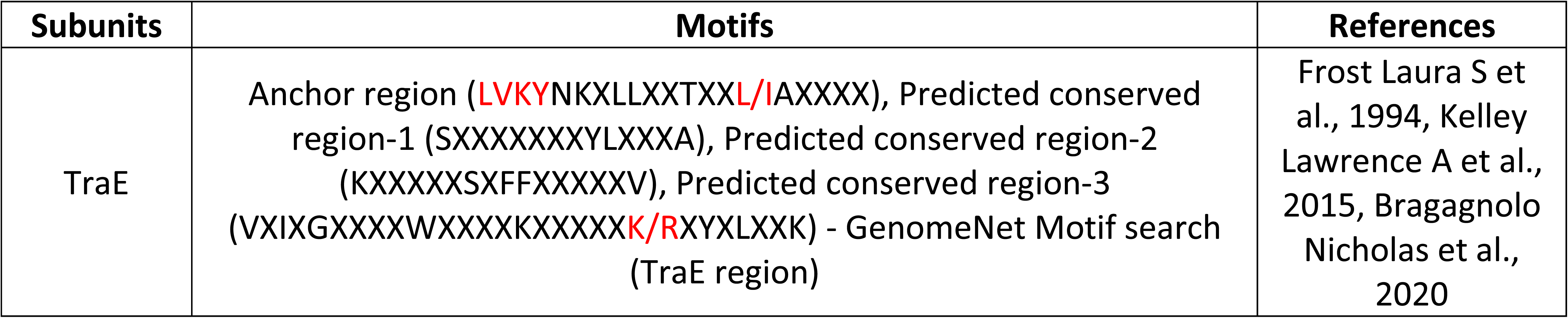
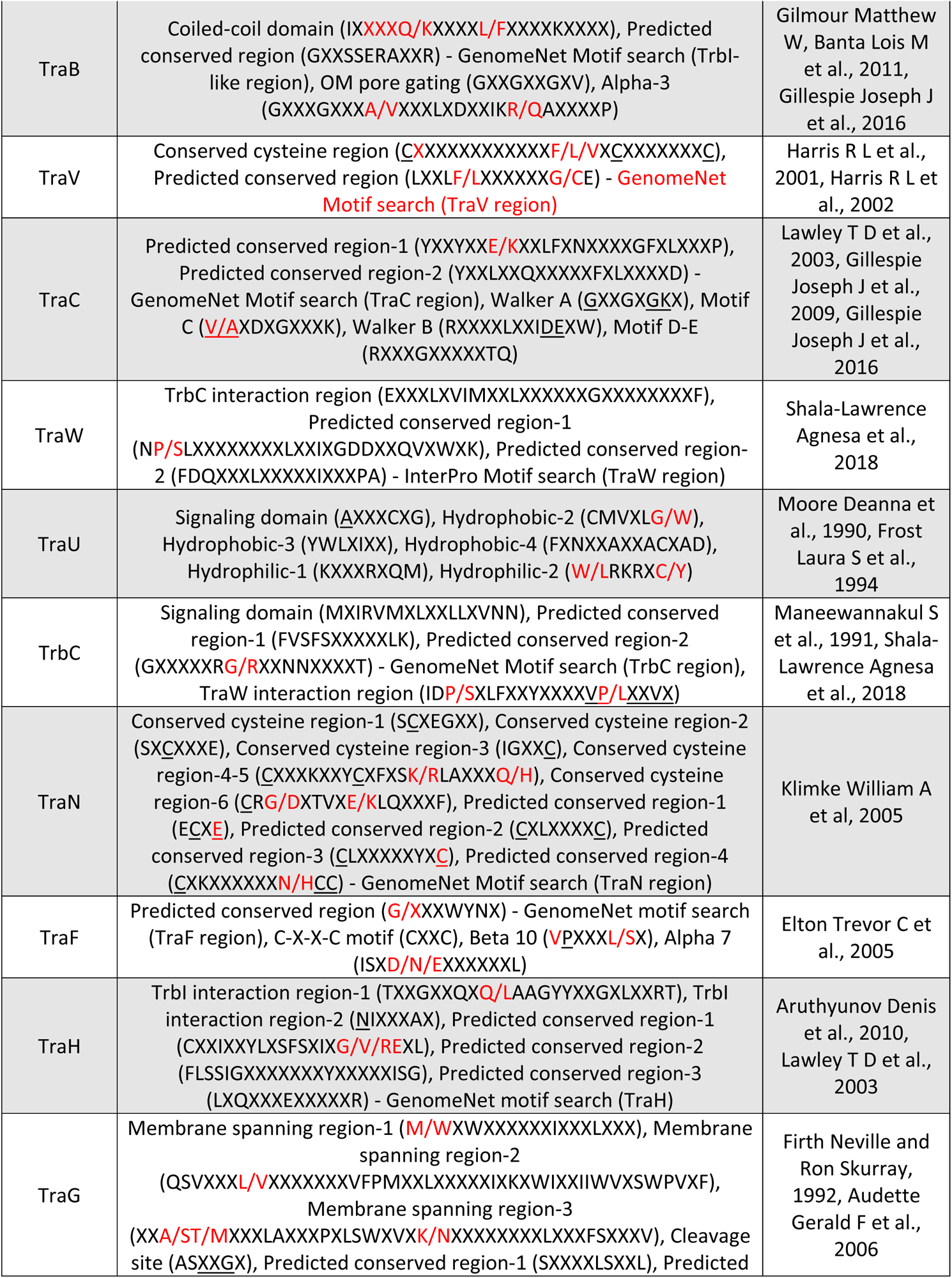
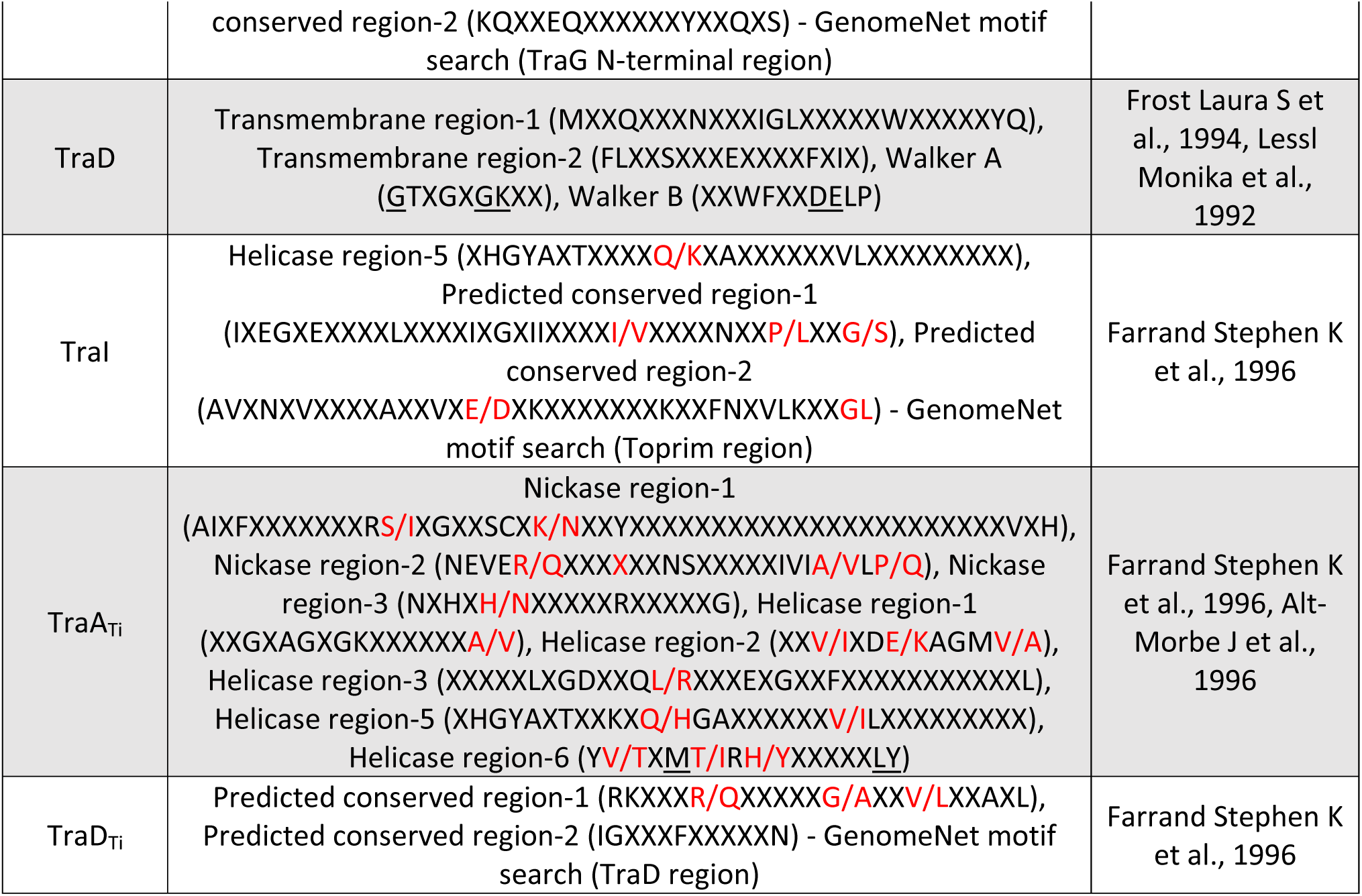
Motifs conserved in Tra and Trb proteins. <u>Underlined</u> base(s) indicate important amino acid in the motif. Red letter indicates variability in amino acid sequence.

Note that TraA_Ti_ found in *Rickettsia* is fragmented into TraA_Ti_ and TraI in *Ot*. The longest *tra* and *trb* genes identified in Ikeda were set as references and were then blasted (nucleotide blast using Geneious Prime) to identify their presence across the 8 strains of Ot (Boryong, Ikeda, Karp, Kato, Gilliam, TA686, UT76, UT176). The amino acid sequences of each Tra subunit from the 8 strains of Ot were then aligned (Multiple alignment - Clustal Omega) to verify their motifs. Those amino acid sequences with difference greater than 10% from full length gene in Ikeda or missing motif(s) are considered pseudogene (truncation).

## Data Availability

All data generated by this work is available within the manuscript and supporting information.

## Author Contributions

Project design and supervision (JS); data analysis and figure preparation (SG, CK, JW, HA, JS, JJG); original manuscript writing (JS); manuscript revisions (SG, CK, JW, JJG, JS).

## Acknowledgements and Funding

We thank all the members of the lab in Cambridge and Bangkok for discussions and support. We are also grateful to Liz Batty and Nick Day in MORU for discussions, encouragement, and support. This work was funded by a Royal Society Dorothy Hodgkin Fellowship DH140154 (JS); Swiss National Science Foundation IZSTZ0_193925 (JS, CK); Wellcome Trust Senior Research Fellowship 224277/Z/21/Z (JS), grants from the National Institutes of Health, USA, R21 AI156762 and R21 AI166832 (JJG) and American Heart Association grant 20PRE35210610 (HA).

